# A five-fold expansion of the global RNA virome reveals multiple new clades of RNA bacteriophages

**DOI:** 10.1101/2022.02.15.480533

**Authors:** Uri Neri, Yuri I. Wolf, Simon Roux, Antonio Pedro Camargo, Benjamin Lee, Darius Kazlauskas, I. Min Chen, Natalia Ivanova, Lisa Zeigler Allen, David Paez-Espino, Donald A. Bryant, Devaki Bhaya, RNA Virus Discovery Consortium, Mart Krupovic, Valerian V. Dolja, Nikos C. Kyrpides, Eugene V. Koonin, Uri Gophna

## Abstract

High-throughput RNA sequencing offers unprecedented opportunities to explore the Earth RNA virome. Mining 5,150 diverse metatranscriptomes uncovered >2.5 million RNA viral contigs. Via analysis of the 330k novel RNA-dependent RNA polymerases (RdRP), this expansion corresponds to a five-fold increase of RNA virus diversity. Extended RdRP phylogeny supports monophyly of the five established phyla, reveals two putative new bacteriophage phyla and numerous putative novel classes and orders. The dramatically expanded *Lenarviricota* phylum, consisting of bacterial and related eukaryotic viruses, now accounts for a third of the RNA virome diversity. Identification of CRISPR spacer matches and bacteriolytic proteins suggests that subsets of picobirnaviruses and partitiviruses, previously associated with eukaryotes, infect prokaryotic hosts. Gene content analysis revealed multiple domains previously not found in RNA viruses and implicated in virus-host interactions. This vast collection of new RNA virus genomes provides insights into RNA virus evolution and should become a major resource for RNA virology.

## Introduction

Viruses are obligate genetic parasites that reproduce inside cells of nearly all life forms and are widely regarded as the most numerous biological entities on earth (Bergh et al., 1989; Hendrix et al., 1999; Mushegian, 2020). For more than a century, the paramount importance of many viruses of animals and plants in human and veterinary medicine, as well as agriculture has been translating into a myopic, parochial view of the virus world. However, the last two decades saw a tectonic shift in the study of viruses, resulting primarily from the momentous advances in genome sequencing, metagenomics and metatranscriptomics. In particular, an enormous, previously unsuspected diversity of viruses with double-stranded (ds) and single-stranded (ss) DNA genomes associated with bacteria, archaea and unicellular eukaryotes has been identified (Call et al., 2021; Dolja and Koonin, 2018; Koonin et al., 2020; Kristensen et al., 2010; Paez-Espino et al., 2016; Roux et al., 2020). Mining metagenomic data has led to major discoveries, such as one of the most abundant groups of bacteriophages in the human gut, the crAss-like phages (Dutilh et al., 2014; Yutin et al., 2021). Undoubtedly, metagenomic data will form the bedrock of our collective, ever-increasing understanding of the critical roles of DNA viruses in shaping microbial diversity and the earth biogeochemistry, including a substantial, direct effect on nutrient cycles on the global scale (Kristensen et al., 2010; Rohwer and Thurber, 2009; Steward et al., 2013; Suttle, 2007). Recognizing the leading role of metagenomics in new virus discovery, the International Committee for Taxonomy of Viruses (ICTV) has adopted a new policy allowing formal recognition of virus species and higher taxa solely on the basis of metagenomic sequence analysis (Simmonds et al., 2017).

In contrast, the role of RNA viruses in diverse microbial ecosystems remained poorly understood. Whereas metagenomic analyses expanded the DNA virome by orders of magnitude, the diversity of the RNA virus world remained disproportionately unexplored, being doubled only twice in recent years. In 2016, pioneering work by Krishnamurthy *et al*. paved the way to harnessing metatranscriptomics (bulk sequencing and analyses of RNA from entire microbial communities) for RNA virus discovery (Krishnamurthy et al., 2016; Zeigler Allen et al., 2017). Since then, several large-scale (meta)transcriptome surveys have uncovered massive amounts of previously undetected, highly diverse RNA viruses (Dolja and Koonin, 2018; Shi et al., 2016). In particular, the study of Shi *et al*. on invertebrate transcriptomes resulted in doubling the number of known RNA viruses (Shi et al., 2016), whereas the recent work of Wolf *et al*. increased the known diversity of RNA viruses a further two-fold by analysis of the RNA sequences in the metavirome (the entirety of RNA sequences from the fraction containing virus-sized entities) from a single, rich habitat, implying a vast, barely sampled global RNA virome (Wolf et al., 2020). Other recent examples of deep forays into RNA viromes include the analysis of fungal transcriptomes (Sutela et al., 2020), as well as metatranscriptomes from different types of soil (Starr et al., 2019; Stough et al., 2018; Wu et al., 2021), and expansion of the RNA phageome of aquatic environments (Callanan et al., 2020).

All RNA viruses share a single hallmark gene that encodes the RNA-dependent RNA polymerase (RdRP), the enzyme responsible for the RNA genome replication (Koonin et al., 2020). Therefore, global analysis of the diversity and evolution of RNA viruses hinges primarily on the detection and phylogenetic analysis of RdRP sequences. Although due to the extreme sequence divergence of the RdRPs, the confidence in the deepest branching in the phylogenetic tree is low, previous analysis identified five well-separated, major clades (Wolf et al., 2018). These have since been formally recognized by the ICTV as phyla within the kingdom *Orthornavirae*, which encompasses all RNA viruses and belongs to the realm *Riboviria*, together with the second kingdom *Pararnavirae* (reverse-transcribing viruses) (International Committee on Taxonomy of Viruses Executive Committee, 2020; Koonin et al., 2020). These five phyla remained robust to the previous doubling of the RNA virus diversity (Wolf et al., 2020), suggesting that the large-scale structure of the RNA virosphere was approaching completeness.

A key requirement for a robust characterization of RNA virus diversity that would provide for a stable taxonomy, at least, at the deepest level, is an adequate sampling of different branches of the RdRP tree. The importance of taxon sampling for obtaining reliable phylogeny is well recognized in molecular phylogenetics (Gouy et al., 2015; Pick et al., 2010; Rodríguez-Ezpeleta et al., 2007), and in the case of RNA viruses, the problem is exacerbated by the fast evolution of RNA genomes that makes individual sequences (singletons) as well as small clusters of related sequences particularly noisy in phylogenetic analyses. Distributions of clusters of homologous sequences, at most levels of clustering, generally follow a power law, with numerous small clusters including singletons and a long tail of larger clusters (Koonin et al., 2002). The larger the clusters are at a given phylogenetic depth, the more information is available to separate cluster-specific features from spurious ones. Small, highly divergent clusters cannot be reliably mapped to the overall phylogenetic framework and thus should not be used to define new taxa. Therefore, a crucial task that is attainable only through large-scale virus diversity surveys is to populate poorly sampled groups of viruses with diverse, representative members. Evidently, such surveys produce a new, large set of singletons and small clusters that await further expansion of sampling (Wolf et al., 2020). Thus, an extensive census of RNA virus genomes from a broad variety of environments and metatranscriptomes is indispensable for the comprehensive characterization of the global RNA virome and better understanding of RNA virus evolution.

Here, we present an analysis of 5,150 metatranscriptomes that resulted in an expansion of the global RNA virome from the previously known 13,282 to 124,873 distinct clusters of virus genome segments at the granularity between the species and the genus level (90% Average Nucleotide Identity, ANI), including two candidate new phyla, 74 new classes compared to the previously established 19 and numerous new orders and families. A key result of this analysis is the major expansion of the parts of the global RNA virome assigned to prokaryotic hosts. Metatranscriptome analysis on this enormous scale necessitated the development of a dedicated computational pipeline for virus discovery and comprehensive phylogenetic analysis that is explicitly linked to the expanded virus taxonomy and is scalable for further, even more expansive studies. The vast collection of RNA virus sequences presented here is expected to become an essential resource for RNA virology.

## Results

### Identification of RNA viruses from diverse metatranscriptomes

To ensure robust and sensitive detection of sequences originating from RNA virus genomes, we devised a computational pipeline that is suitable for analysis of thousands of metatranscriptomes (Figure 1). Because RNA viruses are not expected to be found in DNA sequencing efforts, the pipeline starts by filtering out sequences likely originating from DNA genomes by comparing the metatranscriptomic contigs to DNA sequences from a diverse set of genomes and metagenomes, using a succession of search tools and prefiltering steps, coupled with semi-automatic performance evaluations (see Methods). The remaining contigs (less than 1% of the initial sequence set) are treated as potential RNA virus candidates and are then searched for the presence of the only hallmark protein that is conserved across the kingdom *Orthornavirae*, the RdRP. Briefly, the substantially reduced sequence space following the subtraction of DNA-matching sequences allowed us to perform an exhaustive RdRP scan via an iterative search procedure, using end-to-end six frame translations. This approach was adopted to enable the recovery of RdRPs encoded with non-standard genetic codes and/or programmed frameshifts, combining multiple search tools using Position-Specific Scoring Matrices (PSSMs), Hidden Markov Models (HMMs) or single sequences as queries, and automatically curating the results before using them as seeds for additional searches (see Methods).

**Figure 1.**
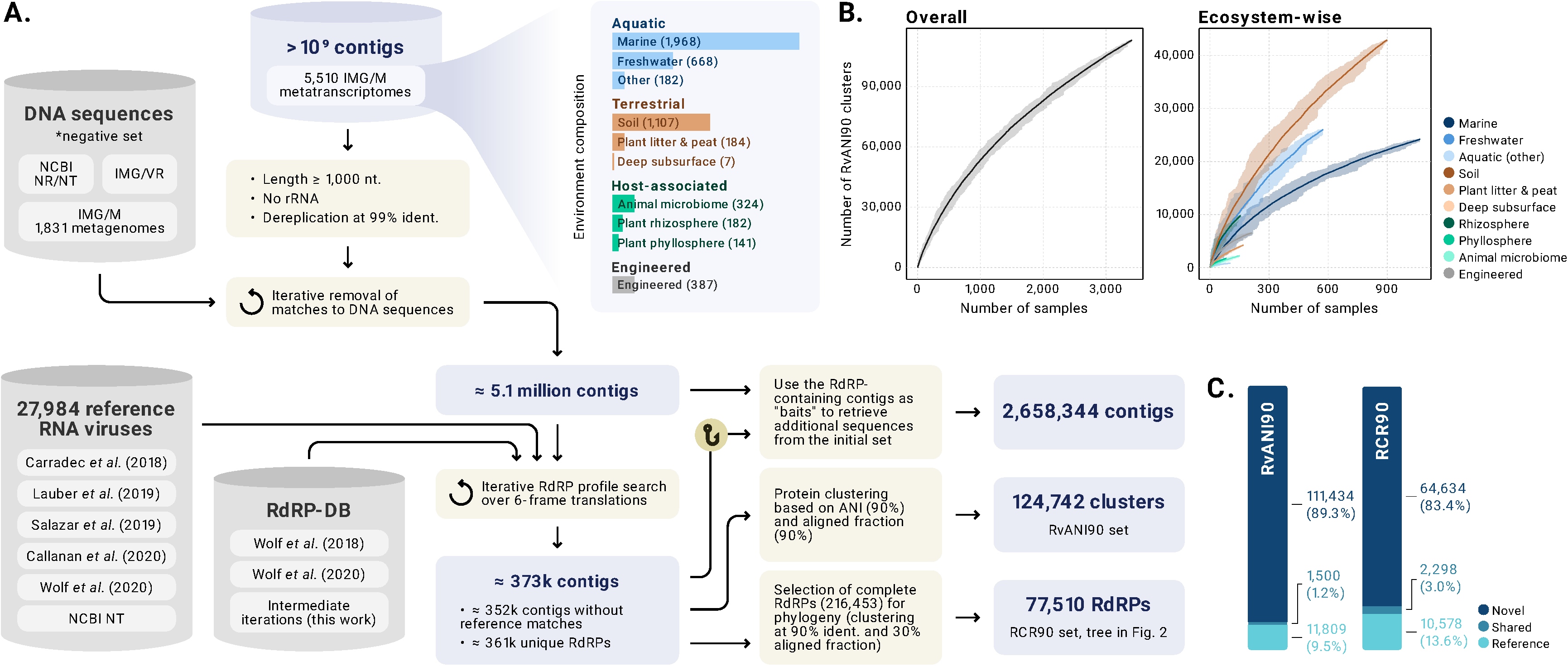
RNA virus discovery pipeline. Flowchart visualisation of the project. **A.** RNA virus discovery pipeline. **B.** RvANI90 Rarefaction curves - accumulation of unique clusters as an outcome of the number of measured biological samples (GOLD field - ITS.PIDs). These values were obtained via bootstrapping; The semi-opaque segments represent the range of measured unique RvANI90 clusters across 25 random subsampling. The central line represents the average of the 25 random samples. Colours indicate the environment type (right chart). **C.** Number of RCR90 clusters (left) and RvANI90 (right), whose members are either entirely “Reference” (contains contigs only from the “Reference set”), “Novel” (only identified in the analysed metatranscriptomes) and “Shared” (contains at least one member of each type).

Among the 5,150 metatranscriptomes queried, 3,598 contained one or more contigs encoding the RdRP core domain of sufficient completeness (presence of motifs A-D or 75% coverage of existing RdRP models) for downstream analyses. We then used the RdRP-encoding contigs as baits to identify additional metatranscriptomic contigs sharing high sequence similarity with the RdRP-encoding ones (including outside the RdRP core; see Methods). Altogether 2,658,344 RNA virus contigs were identified and supplemented with 27,984 sequences from published sources for the purpose of novelty estimation (Fig. 1A). Of these, 348,762 contigs represented a deduplicated, non-redundant sequence set of length ≥1kbp. These sequences were grouped into 124,743 clusters sharing 90% average nucleotide identity (RNA virus ANI90 clusters or RvANI90, see Methods), of which only 13,308 (10.7%) contain at least one sequence that was previously known. Thus, our metatranscriptome mining approach resulted in a roughly ∼9-fold expansion of the global RNA virome, at the ANI90 level of diversity.

As expected, the RNA virus sequence clusters displayed a power law-like distribution by size, which is dominated by small clusters, with a long tail of large ones, the largest cluster including 429 contigs (Suppl. Fig. S1). Based on the accumulation curve, the global diversity of RNA viruses evaluated at the RvANI90 cluster level showed no sign of saturation (Fig. 1B), with a particularly high richness in soil environments (Fig. 1B). About 5.8% of the RdRP encoding contig set showed evidence of utilising alternative genetic codes (green arcs in Fig. 2), and about 0.5% showed shuffling of the order of motifs (known as “domain permutation”) within the RdRP coding sequence (orange arcs in Fig. 2). These atypical sequences are usually missed by standard RdRP sequence searches, but did not escape our 6-frame search approach, and are mostly concentrated in a handful of specific clades (see below).

**Figure 2.**
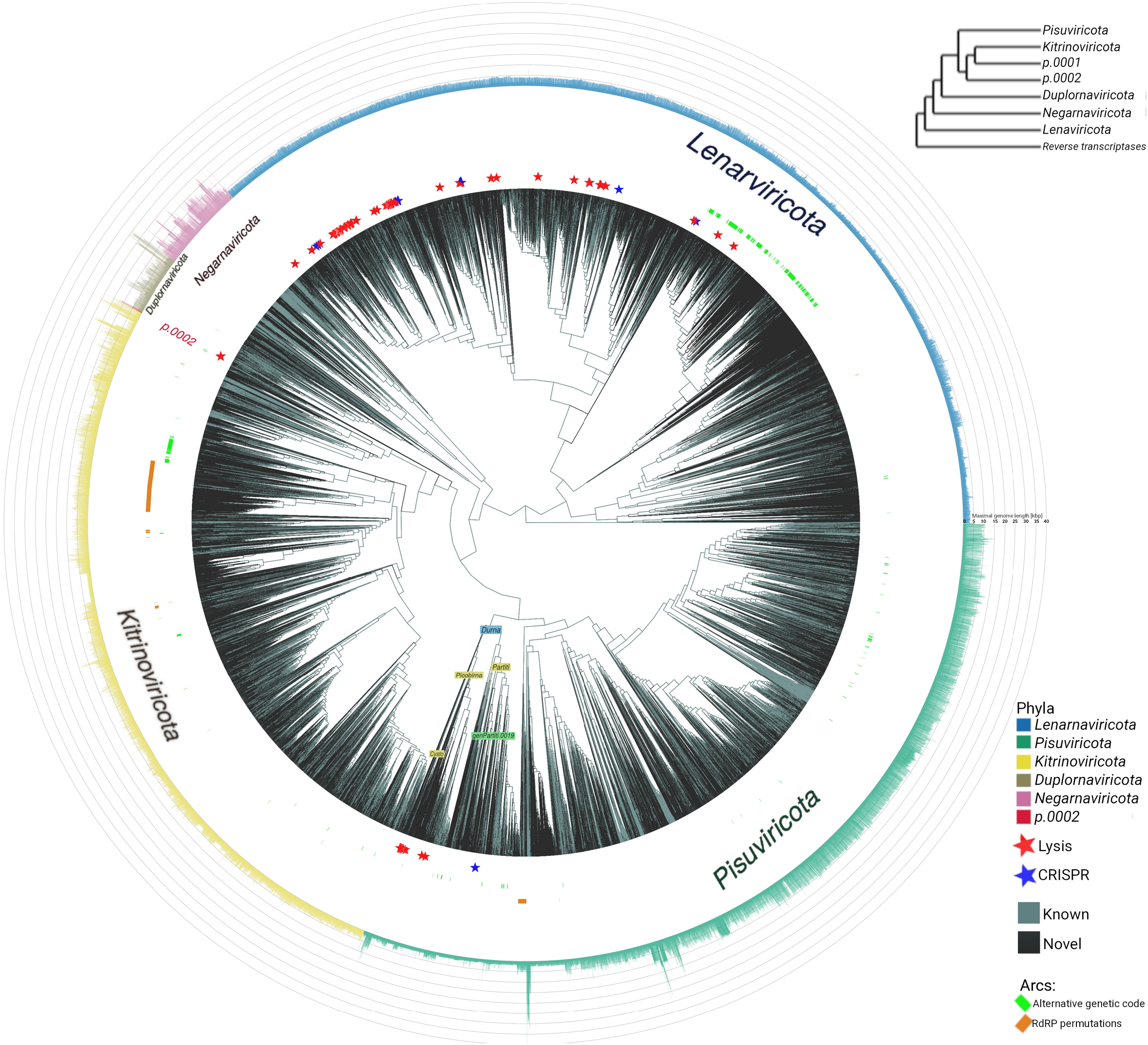
Phylogenetic reconstruction of the global RNA virosphere. An ultrameterized circular tree rooted using reverse transcriptases as an outgroup. Visualised programmatically using the ggtree and ggtreeExtra R libraries (Xu et al., 2021; Yu et al., 2018). Branches are colored Black, unless any of their descendants contains at least 1 sequence from the “Reference set” (Cyan). Tips aligned with stars indicate evidence of prokaryotic host - CRISPR spacer match in Blue, and bacteriolytic domain in Red. Green arcs indicate clades where ≥50% of the associated sequences carry evidence of alternative genetic code usage. Orange arcs indicate clades where ≥50% of the RdRPs show evidence of motif permutation. The 5 previously established phyla, and the proposed candidate phylum p.0002 are colour coded in both the text and the bar-plot in the outermost ring, which represents the maximal contig/genome length observed for each RCR90 cluster (i.e. tree tip). Some key taxa are text labelled directly on the tree. A fully taxonomically labelled version of this figure in high resolution is available in the project’s Zenodo repository (see code and data availability).

### RdRP-based RNA virus phylogeny and major diversity expansion across taxonomic ranks

To build a global RNA virus phylogeny, we first extracted the full-length RdRP core domain sequences (operationally defined as covering motifs A–C) and clustered them at the 90% amino acid identity threshold, arriving at 77,510 representatives (RCR90 set, for RdRP Core Representatives at 90% identity, see Methods). Even when reduced to the RCR90 granularity, the dataset remained too large and too diverse to be directly amenable for multiple sequence alignment construction and subsequent phylogenetic analysis with advanced maximum likelihood phylogenetic methods, such as IQ-tree. Therefore, we used an iterative clustering procedure whereby sequences were initially clustered with MMSeqs2 and cluster alignments were produced using MUSCLE5; then, the clusters were compared using HHSEARCH, further clustered using cluster-to-cluster scores and aligned to each other using HHALIGN; finally, the cluster consensus sequences were aligned using MUSCLE5.

The resulting RdRP tree comprised 77,520 representative sequences (77,510 RCR90 sequences and 10 representative reverse transcriptases (RT) included as an outgroup; Fig. 2). Even with this dramatic sequence expansion, the five phyla previously established (Wolf et al., 2018; Wolf et al., 2020) remained largely monophyletic. In addition, the tree included two groups below the base of the phylum *Kitrinoviricota*, which were further analysed in greater detail (see below).

The monophyly of the major branches in the RdRP tree, in particular the previously established five phyla, was verified by subsampling. Briefly, representatives of virus families were repeatedly randomly sampled, phylogeny was reconstructed from the multiple alignment of each sample, the positions of the phyla clades were traced, and the quantitative measure of their monophyly was calculated (see Methods). In most of the samples, the five previously established phyla stayed largely monophyletic (Suppl. Fig S2a). Sequences that tended to break the phylum-level monophyly formed a sharply biased subset, with flaviviruses (class *Flasuviricetes*) being by far the most common “offenders”. Our current analysis of the full tree placed *Flasuviricetes* inside *Pisuviricota*, whereas in the previous analyses this class was the basal clade of *Kitrinoviricota*. The instability of the placement of flaviviruses in subsampled trees is another indication of the uncertainty of their phylogenetic position. The families *Reoviridae, Picobirnaviridae, Cystoviridae* and several new families also often broke away from their respective phyla although the consensus tree placed *Picobirnaviridae* and *Cystoviridae* confidently inside *Pisuviricota* (see below).

When the subsampled trees were reduced to the lowest common ancestor of each of the five phyla, the deepest branching order was found to be robust, with *Pisuviricota* and *Kitrinoviricota* forming the crown group in the consensus tree, and *Lenarviricota* and *Negarnaviricota* occupying the basal position (Fig. 2, top right inset). Recapitulating previous analyses (Wolf et al., 2018; Wolf et al., 2020), when the tree was rooted by the RT, distant homologs of the RdRPs, the deepest branch within the kingdom *Orthornavirae* was the phylum *Lenarviricota* that includes leviviruses (positive-sense RNA bacteriophages; class *Allassoviricetes*) and their apparent direct descendants among the viruses of eukaryotes, namely, mitoviruses (*Howeltoviricetes*; reproducing in mitochondria), narnaviruses (*Amabilivirecetes*) and botourmiaviruses (*Miaviricetes*). Although supporting this branching order beyond reasonable doubt might not be feasible, this position of *Lenarviricota* is biologically plausible, placing the origin of *Orthornavirae* in the bacterial domain. In contrast, the placement of *Negarnaviricota* as the next deepest branch in the tree was unexpected given that negative sense RNA viruses have been isolated almost exclusively from animals and plants. Potentially, this position of *Negarnaviricota* might reflect an ancient origin, suggesting existence of yet unidentified negative-sense RNA viruses infecting prokaryotes. More likely, however, this is an artefact of phylogenetic analysis, perhaps caused by acceleration of evolution at the base of *Negarnaviricota*.

Comparison of the phylogenetic depths of the present RdRP phylogeny and the previously reported trees reflected the expansion of the global RNA virome as measured by the total-branch-length (TBL), which increased roughly 5 fold compared to the previous analysis (Wolf et al., 2020). To convert the RdRP phylogeny into a tentative taxonomic scheme, we developed a semi-quantitative approach for assigning taxonomic ranks to unclassified nodes in the global RNA phylogeny based on neighbouring well-established taxa (see Methods for details). The new taxa were designated according to the rank and prefixed by *p*, *c*, *o, f* and *g* for phylum, class, order, family and genus, respectively, followed by an ordinal number for new taxa of that rank. Optionally, taxa that are associated with a previously described taxon were terminated with a label “base”, e.g., *f.0127.base-Noda* is the 127th new family that is basal to *Nodaviridae* in the RdRP tree (Table S1).

Application of this approach resulted in a roughly five-fold expansion of the RNA virus diversity at all ranks below the phylum, compared to the results of the latest RNA virome analysis (Wolf et al., 2020), and an order of magnitude increase compared to the set analysed in 2018 (Wolf et al., 2018) (Table 1). When broken down by phyla, the largest expansion at all taxonomic ranks was observed within the phylum *Lenarviricota*, followed by *Kitrinoviricota*, and then, *Pisuviricota*. By contrast, the phyla *Duplornaviricota* (double-stranded RNA viruses) and *Negarnaviricota* (negative-strand RNA viruses) remained largely stable, with only a few taxa added (Fig. 2; Table S1).

**Table 1.** Expansion of the global RNA virome resulting from metatranscriptome analysis

| Virus taxon rank | Number of known taxa | Updated number of taxa | Fold increase |
| --- | --- | --- | --- |
| RvANI90 cluster | 13,282 | 124,873 | 9.4 |
| RCR90 cluster | 12,862 | 77,510 | 6.0 |
| Family | 98 | 489 | 4.9 |
| Order | 26 | 121 | 4.7 |
| Class | 19 | 93 | 4.9 |
| Phylum | 5 | 7 | 1.4 |

In addition to this major expansion of the virus taxa reflected in the RdRP phylogenetic tree, some of the novel RNA viruses identified via the RdRP-based profile searches were semi-automatically discarded from the subsequent phylogenetic analysis because the boundaries and some of the motifs of the core RdRP domain could not be reliably identified. In total about 39,000 contigs that formed 24,742 RvANI90 clusters that contained a confidently identifiable RdRP signature, were excluded through this quality control step.

### Putative new phyla and classes of RNA viruses

Two of the most divergent new clades were positioned below the base of *Kitrinoviricota* in the RdRP phylogeny and, in principle, could be included in an expanded version of this phylum. The first of these branches (termed *p.0001*) included mere 3 RCR90 clusters, and therefore, its further analysis was deemed premature. The second of these deep branches possessed unique features that appeared better compatible with the establishment of a new phylum over the expansion of *Kitrinoviricota*. This candidate phylum (*p.0002* for short) consisted of 234 contigs from 31 RvANI90 clusters, of which the most complete specimens had an average length of about 12kb and encompassed 10 ORFs. Most of the predicted proteins, except for the RdRP, showed no significant sequence similarity to any available protein sequences. A notable exception, however, was a gene in one of the two tentative families comprising this putative phylum that encoded a protein of either the M15 or M35 family of zinc metallopeptidases implicated in cell lysis (see below). The ORFs in the genomes of the *p.0002* viruses are tightly spaced, and examination of the nucleotide sequences upstream of the coding regions showed the recurring presence of ribosome-binding (Shine Dalgarno, SD) motifs used in prokaryotic translation (Fig. 3A). Taken together, these observations strongly suggest that *p.0002* is markedly different from the members of the phylum *Kitrinoviricota* that are known to infect eukaryotes. Most likely, *p.0002* represents a new phylum of RNA viruses infecting prokaryotes, as discussed below, although some caution is due with this interpretation because at present *p.0002* includes only 30 RCR90 leaves.

**Figure 3.**
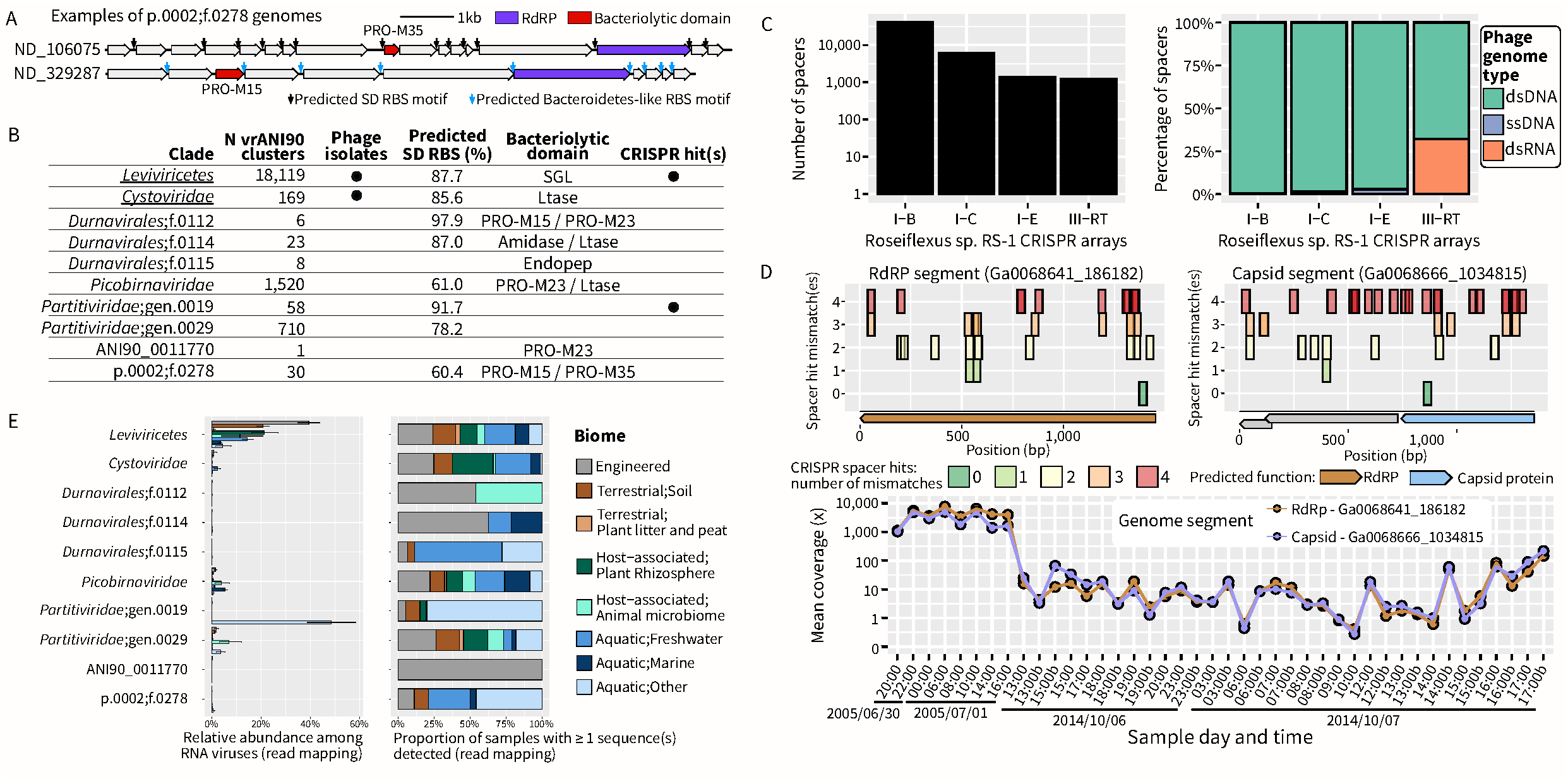
Diversity and abundance of known and predicted prokaryotic RNA viruses. **A.** Genome map of two distinct contigs assigned to the proposed new phylum p.0002 and proposed new family f.0278. Predicted cds are colored based on functional annotation, and predicted RBS motifs are indicated with colored arrows. **B.** Overview of recognized (underlined) and predicted prokaryotic RNA viruses. For each group, the type of evidence supporting its association with prokaryotic hosts is indicated. For Shine-Dalgarno (SD) RBS, clades were considered as likely composed primarily of phages if ≥ 50% of the predicted cds were associated with an SD motif, after excluding genes predicted on the edges of contigs. Ltase: Lytic transglycosylase, lysozyme superfamily fold, SGL: “Single-gene lysis” (cell wall biosynthetic inhibitors), PRO-M15: Zn-DD - carboxypeptidase (sensu PF08291.13) – Endolysin in dsDNA and some ssDNA viruses, PRO-M35: M35 peptidase; zinc metalloendopeptidase family M35 (MEROPS), PRO-M23: M23-family metallopeptidases, Amidase: N-acetylmuramoyl-L-alanine amidase, Endopep: L-alanyl-D-glutamate endopeptidase. **C.** CRISPR spacer landscape of *Roseiflexus* sp. RS-1 in Yellowstone hot springs, including spacers matching newly-identified *genPartiti.0019* phages. The left panel displays the total number of spacers identified for each of the 4 types of CRISPR arrays encoded in the genome of *Roseiflexus* sp. RS-1 (see Fig. S3). The right panel presents the type of phage (dsDNA, ssDNA, or RNA) for which hits to CRISPR spacers were identified for each CRISPR type. **D.** Example of a predicted pair of RdRP- and capsid-encoding segments from a newly-identified *genPartiti.0019* phage. Top panel: hits to CRISPR spacer matches are indicated alongside a genomic map for each segment. The number of mismatches is indicated on the y-axis, while the position of the hit is indicated on the x-axis. The bottom panel displays the relative abundance of both segments across a metatranscriptome time-series. **E.** Relative abundance of different groups of (recognized or predicted) prokaryotic RNA viruses across biomes. Here, only datasets dominated by prokaryotic sequences (“P-dominated”) containing at least 10 prokaryotic RNA viruses were considered. The right panel presents a breakdown of the biome distribution for each clade, calculated from a balanced dataset composed of random subsamples of 50 samples per environment. This random subsampling was performed 100 times, and the average values are plotted here.

Another highly divergent virus group and a putative candidate phylum was RvANI90_0011770, one of the clusters whose RdRP were omitted from the tree in Fig. 2, because they distorted the RdRP alignment (hence no *p* designation). The largest of the 55 contigs comprising this group were 10-12 kb long and encompassed 7 to 9 closely spaced ORFs albeit with no conserved SD motifs. One of these ORFs encoded the highly divergent RdRP and another one encoded a predicted lysis enzyme (see below), whereas the remaining ORFs showed no similarity to any known proteins. Notably, these contigs all originate from 27 different active sludge samples. With the same caveats as pointed out for *p.0002*, RvANI90_0011770 might represent a novel phylum consisting of previously unknown RNA bacteriophages.

Within each phylum, we observed a substantial increase in the class-level diversity (assessed quantitatively as outlined in the Methods) of the four established phyla which included: 14 new classes over the 4 previously known in *Lenarviricota*, 18 new classes over the 4 previously known in *Pisuviricota*, 20 new classes versus 3 previously known in *Kitrinoviricota,* and 18 new classes over 6 previously known in *Negarnaviricota*. Only in *Duplornaviricota* was there parity, with two new class-level clades identified in addition to the two recognized classes. Overall, according to our analysis, the five established phyla of *Orthornavirae* contained 91 classes compared to the 19 previously established, and 391 new families to be added to the previously recognized 98 (Table 1; Table S1).

### Major expansion of the range of RNA viruses associated with bacteria

So far, the overwhelming majority of RNA viruses have been associated with eukaryotic hosts, with only two groups known to infect bacteria, namely, leviviruses (*Leviviricetes* within the phylum *Lenarviricota*) and cystoviruses (*Vidaverviricetes* here reassigned to *Pisuviricota* as described below). In the preceding section, we describe two putative new phyla, *p.0002* and RvANI90_0011770, that consist of viruses with likely prokaryotic hosts. Prior to the recent advances in virus metagenomics, both leviviruses and, particularly, cystoviruses, included small numbers of viruses with narrow host ranges. In this work, an expansion of *Cystoviridae* diversity was evident as this family was represented by 8 RCR90 sequences from published sources and 132 new RCR90 sequences from metatranscriptomes. Although the diversity of the leviviruses was substantially expanded in recent studies (Callanan et al., 2020), this trend persisted in the present metatranscriptome analysis, with 13,512 new RCR90 leaves of *Leviviricetes* compared to the 1,940 previously known.

The expanded phylum *Lenarviricota* now accounts for more than a third of the RNA virus RCR90 sequences and includes the four largest families of RNA viruses (Fig. 2; Table S1). The top and fourth largest families, *Steitzviridae* and *Fiersviridae*, respectively, are bona fide *Leviviricetes* bacteriophages. The second-largest family, *Botourmiaviridae*, consists of eukaryotic viruses that appear to have evolved from a common ancestor with *Leviviricetes* RNA phages, with the capsid-less *Narnaviridae* and *Mitoviridae* (the latter comprising the third largest family of RNA viruses) as intermediates (Dolja and Koonin, 2018; Koonin et al., 2020).

In addition to the major expansion of the bacteriophage taxa within *Lenarviricota*, converging lines of evidence suggest reassignment to bacterial hosts for several groups of viruses previously thought to solely infect eukaryotes as well as inference of prokaryotic hosts for several new groups (Fig. 3B). These indications include the detection of bacteriophage-specific lysis proteins, similarity to CRISPR spacers, and presence of SD motifs, used in bacteria and archaea as ribosome binding sites, upstream of predicted genes (see Methods). Viruses of prokaryotes now also appear to be interspersed with those infecting eukaryotes within *Pisuviricota*. Specifically, the family *Cystoviridae*, which migrated from *Duplornaviricota* to *Pisuviricota* in the current RdRP phylogeny, now forms a strongly supported branch with picobirnaviruses and partitiviruses, two groups of double-stranded RNA viruses embedded in the midst of the positive-sense RNA viruses comprising this phylum (Fig. 2). Within this *Durnavirales* order, several clades showed unexpected conservation of SD motifs in the 5’ untranslated regions (UTRs), suggesting that these viruses infect bacteria, many of which employ SD motifs for translation initiation (Bahiri Elitzur et al., 2021; Hockenberry et al., 2018). These putative new RNA phages include members of *Picobirnaviridae*, for which the presence of SD motifs was previously noted (Boros et al., 2018; Wang and Krishnamurthy, 2018), along with two deep-branching families (*f.0114.base-Cysto* and *f.0112.base-Cysto)* and two new genera within the partitivirus clade (namely, genPartiti.0029, genPartiti.0019.base-Deltapartitivirus) (Table S2, Fig. 3B).

Another, independent line of evidence of the bacterial association of most of the novel RNA viruses is the conserved occurrence of bacteriolytic proteins (Fig. 3B). Numerous dsDNA bacteriophages and dsRNA viruses of the *Cystoviridae* family encode one or more lytic enzymes, known as endolysins, that possess diverse enzymatic activities and degrade bacterial peptidoglycan (Cahill and Young, 2019; Criel et al., 2021; Vázquez et al., 2021). In contrast, leviviruses induce host cell lysis by inhibiting the peptidoglycan biosynthesis via small membrane proteins that comprise the so-called single gene lysis (Sgl) system (Cahill and Young, 2019). The *sgl* genes typically overlap or are completely nested within other levivirus genes, such as RdRP (Chamakura and Young, 2020). To identify lysis proteins that could be encoded in newly discovered RNA virus genomes, we employed a collection of domain profiles for known bacterial and phage lysis proteins (Fig. 3B). This custom domain search yielded ∼546 significant matches to lysis protein profiles (see Methods). As would be expected, the majority of the predicted lysis proteins were identified in genomes of the *Leviviricetes* (469), and *Cystoviridae* (17).

Members of the *Leviviricetes* invariably encoded Sgl proteins, typically, nested within the *RdRP* gene. Cystoviruses are known to encode lytic transglycosylases with the lysozyme superfamily fold that digest peptidoglycan during both virion entry and egress (Dessau et al., 2012; Mindich and Lehman, 1979). By contrast, the expanded set of cystoviruses discovered here encode a diverse repertoire of peptidoglycan-digesting enzymes. In agreement with previous observations, all members of the *Cystoviridae* clade *sensu stricto* were found to encode lytic transglycosylases, whereas viruses of the *f.0114.base-Cysto* clade encode either lytic transglycosylases or N-acetylmuramoyl-L-alanine amidases. All members of the *f.0112.base-Cysto* clade encode metallopeptidases of the M15 or M23 families (Table S2) that both cleave bonds within the cross-linking peptides and are commonly encoded by dsDNA phages (Oliveira et al., 2013). In addition, certain members of the *f.0112.base-Cysto* clade encode lipases that also could facilitate host cell lysis. Finally, the *f.0115.base-Cysto* members encode a putative L-alanyl-D-glutamate endopeptidase, another enzyme that commonly functions as an endolysin in dsDNA phages (Cahill and Young, 2019; Oliveira et al., 2013). This clade-specific distribution of endolysin genes in cystoviruses indicates that, as in the case of dsDNA bacteriophages, lysis genes are subject to relatively frequent non-homologous replacement, potentially linked to adaptation to new hosts.

Two other groups of RNA viruses were found to encode lysis proteins, namely, picobirnaviruses and viruses of the putative family *f.0278* in the proposed new phylum *p.0002*. Six picobirnaviruses encode either lytic transglycosylases homologous to phage lambda lysozyme or M23-family metallopeptidases homologous to those encoded by cystoviruses mentioned above. Members of the proposed family *f.0278* encode either M15 or M35 family zinc metallopeptidases (Table S2). The M15 family enzymes mediate host cell lysis during dsDNA phage infection, as in the case of phage T5 (Kutyshenko et al., 2021) and were predicted also to function as lysis proteins of some ssDNA bacteriophages (Roux et al., 2012), whereas M35-family enzymes have not been previously linked to phage egress. However, given that the two enzymes are mutually exclusive in *f.0278* viruses and the corresponding genes occupy equivalent positions, we propose that both M15 and M35 family proteins function as endolysins. The conservation of M15 and M35 family proteins in *f.0278* strongly supports bacterial host assignment for this virus family. RvANI90_0011770, a putative new phylum of RNA bacteriophages identified in RdRP profile searches but not included in the main tree (see above), showed similar conservation of an M23-family metallopeptidases.

The final line of evidence for virus assignment to a bacterial or archaeal host was the detection of significant sequence similarity between RNA viruses and CRISPR spacers, that is, short viral sequence segments integrated into bacterial and archaeal CRISPR arrays and involved in adaptive immunity (Makarova et al., 2020; Mohanraju et al., 2016). Although most known CRISPR systems target DNA templates, and so far only a single match from a CRISPR spacer to an RNA bacteriophage has been reported (Wolf et al., 2020), a large subset of type III CRISPR systems encodes a reverse-transcriptase (RT), and both type III and type VI CRISPR systems can protect bacteria against RNA bacteriophages in laboratory experiments (Abudayyeh et al., 2016; Makarova et al., 2020; Silas et al., 2017). Here, we compared the entire collection of newly identified RNA virus genomes to the IMG database of ≥ 50 million spacers, and after removal of unreliable hits and/or taxonomically inconsistent connections (e.g., a single CRISPR spacer hit to a sequence belonging to a clade of otherwise eukaryotic viruses), curated CRISPR spacers matches were identified to 161 RNA viruses from 23 RvANI90 clusters, across two clades: *Leviviricetes*, and *genPartiti.0019* (Figs. 3B) (Table S2). All the matches to members of *Leviviricetes* were from the IMG database of metagenome-derived CRISPR spacers, mostly from short contigs, for which there was no reliable taxonomic information and no *cas* genes were detectable (Table S3). By contrast, the matches to members of *genPartiti.0019* were specifically associated with populations of *Roseiflexus* sp. RS-1, and were thus further analysed. This filamentous anoxygenic phototrophic bacterium of the phylum *Chloroflexi* is a dominant member of microbial mats in Mushroom Spring (Davison et al., 2016), which is the origin of the *genPartiti.0019* sequences. The genome of *Roseiflexus* sp. RS-1 encodes four distinct CRISPR systems, with one subtype III-B locus encoding a RT fused to the Cas1 protein (van der Meer et al., 2010)(Supplementary Fig. S3).Compiling spacers across 16 metagenomes, each of the CRISPR arrays could be associated with ≈1,000 to ≈40,000 spacers, but spacers matching *genPartiti.0019* sequences were near-exclusively (all but one) detected in the RT-encoding III-B array, suggesting that these were acquired from RNA templates (Fig. 3C). These CRISPR spacer matches were detected across a period of 9 years (2005 – 2014) and showed clear dynamics of spacer gain/loss through time, indicating a likely virus-host association (Supplementary Fig. S3).

Because all *genPartiti.0019* contigs encoded RdRP alone, whereas related partitiviruses have segmented genomes, in which the capsid and other proteins are encoded in separate segments, we reasoned that the same Mushroom Spring metatranscriptome assemblies should also contain contigs coding for the corresponding capsid proteins. Combining matches to spacers from the RT-encoding Type III-B array of *Roseiflexus* sp. RS-1, the absence of corresponding sequences in the Mushroom Springs DNA metagenome, and strong relative abundance correlation (>0.9) to at least one *genPartiti.0019* RdRP-encoding sequence, we identified 88 potential capsid-encoding contigs (Fig. 3D, Supplementary Table S3). Nearly all these candidates (86 of 88) encoded proteins with best alignment to HMM profiles of known partitiviruses capsids (see Methods), at low levels of sequence similarity, but clear similarity of the predicted secondary structures (Supplementary Fig. S3). Taken together, these observations strongly suggest that members of the novel *genPartiti.0019* clade represent a new group of RNA bacteriophages related to partitiviruses and infecting *Roseiflexus* sp. RS-1.

Finally, we evaluated the relative abundance and prevalence of the different groups of (putative) prokaryotic RNA viruses across biomes. Within datasets dominated by prokaryotic hosts (“P dominated”, see below), most of the established or predicted RNA phage clades were detected across a broad range of biomes, but *Leviviricetes* remained by far the most abundant group of prokaryotic RNA viruses, except in some Yellowstone hot springs dominated by the newly-discovered *genPartiti.0019* (Fig. 3E). Taken together with the global diversity analysis, these results signal a substantial shift in our view of the global virome composition, with a sharp increase in the diversity and relative abundance of prokaryotic RNA viruses, challenging the traditional concept that RNA viruses almost exclusively infect eukaryotes (Koonin et al., 2015).

### Differential distribution of RNA virus phyla and classes across samples and habitats

Exploring RNA virus diversity across ∼3,600 metatranscriptomes provides a unique opportunity to evaluate the global distribution of major RNA virus groups. Our global RNA virus catalogue spanned all continents and oceans, reflecting the ubiquity of RNA viruses on Earth (Fig. 4A). Previous metagenomic studies have shown that the distribution of DNA viruses is shaped by the environment type and host community composition (Gregory et al., 2019; Martinez-Hernandez et al., 2017; Roux et al., 2016), and the same factors are likely to determine the RNA virus distribution. An additional factor to consider, however, is the protocol used to generate metatranscriptomes, namely, whether total RNA was sequenced without polyA selection or rRNA-depletion, transcripts were selected using poly(A) enrichment, or ribosomal RNAs were depleted (Gann et al., 2021). Our datasets were processed through all three types of protocols, although they were dominated by rRNA-depleted ones (67% of datasets, Supplementary Fig. S4). We further verified that poly(A)-enriched and total RNA datasets were dominated by eukaryotic sequences, whereas rRNA-depleted datasets consisted mostly of sequences from bacteria and archaea (Fig. S4). The datasets were then separated into three groups: “Eukaryote-dominated” (N=811), “Prokaryote-dominated” (N=2706), and “Mixed” (N=452), based on the taxonomic composition of non-viral contigs. Notably, even Prokaryote-dominated metatranscriptomes frequently included few to no predicted prokaryotic RNA viruses (Fig. S4).

**Figure 4.**
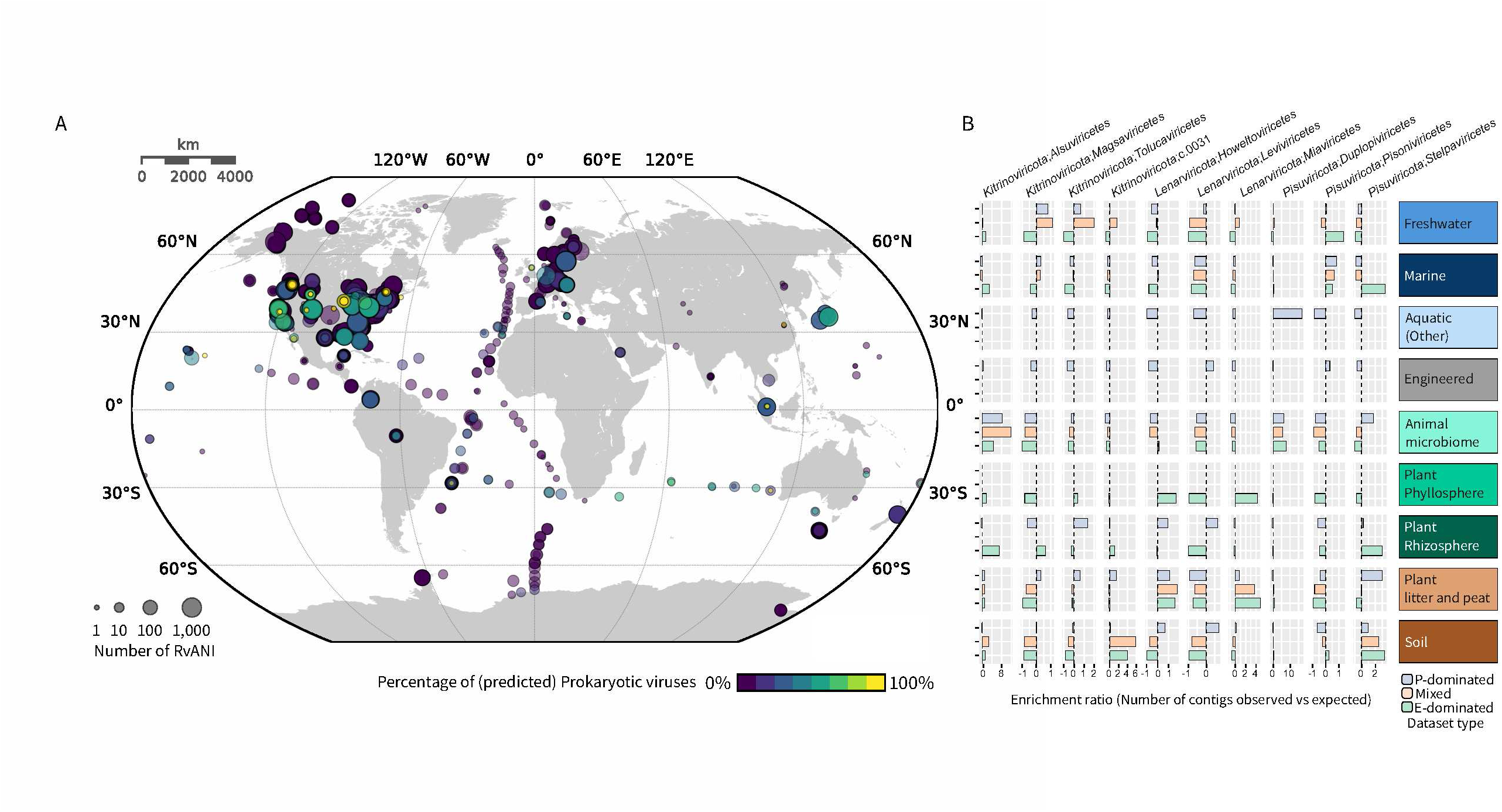
Global distribution of RNA virus sequences. **A.** Map of metatranscriptomes samples from which RNA viruses were identified. For each sample, the circle size reflects the number of distinct RNA viruses (RvANI clusters) identified, and the circle colour indicates the proportion of sequences predicted as prokaryotic viruses. **B.** Relative proportion of RNA virus (proposed) classes (x-axis) detected across ecosystem types (y-axis). To take into account the total number of genomes detected for each class and the total number of samples for each ecosystem type, the counts are represented as enrichments compared to the expected number of genomes for each combination under the hypothesis of even distribution of all classes across all ecosystems. For this analysis, datasets were divided into “E-dominated” (mostly composed of eukaryotic transcripts), “P-dominated” (mostly composed of prokaryotic transcripts), and “Mixed” (see Fig. S4). Enrichments are shown only for combination of ecosystem and dataset type (e.g. “Marine P-dominated”) for which at least 20 metatranscriptomes with at least 1 RNA virus detected were available.

When evaluating the numbers of distinct RNA viruses detected across samples, individual RNA virus classes typically showed clear distribution patterns across dataset types and environments, likely reflecting the distribution of their primary host group (Fig. 4B,E). For instance, sequences from the main class of RNA phages (*Leviviricetes*) were consistently enriched in P-dominated samples from Engineered, Rhizosphere, and Soil environments (Fig. 4B). This is consistent with previous reports of predicted RNA phages representing only a substantial portion of the RNA viral community in some biomes, including marine and freshwater samples, and hints at an uneven ecological distribution of RNA phages at the global scale (Callanan et al., 2020). Conversely, in the same *Lenarviricota* phylum, members of the *Miaviricetes* class, infecting mostly fungi, invertebrates and plants, were associated with E-dominated and Mixed datasets, whereas members of the *Howeltoviricetes* class, which includes mitoviruses, were common in all types of samples but found preferentially in plant-associated environments also rich in fungi. This differential distribution of RNA virus classes can help infer potential host range and ecological niches for different taxa, and represents a roadmap for what type of environment should be further sampled with the goal of comprehensive characterization of the RNA virosphere.

### Modular evolution of RNA virus genomes

Comparative analysis of the expanded RNA virome provided insights into the evolution of RNA virus genomes and their expression strategies. We identified multiple virus groups with pronounced genomic rearrangements, such as fission or fusion of genomic segments (monopartite versus segmented genomes) as well as segmentation and rearrangement of the ORFs encoding polyproteins. For instance, common genomic rearrangements involving the CP-encoding structural module were observed in the order *Picornavirales* and related viruses, where within the same group, capsid proteins are encoded downstream or upstream of the genome replication module, as part of the same polyprotein or as separate proteins (Figure S5. Genome maps). Known viruses of *Benyviridae*, *Picobiranviridae,* and *Botourmiaviridae* have segmented genomes, with the capsid proteins and RdRPs encoded on different segments. By contrast, in several newly identified members of these families, the segments encoding RdRP and CP are fused. Similar rearrangements are known to exist in *Nodaviridae, Secoviridae*, *Closteroviridae* and *Rhabdoviridae* (Walker et al., 2020; Wolf et al., 2020), and were further confirmed and extended in this study. Collectively, these observations emphasise the inherent plasticity of RNA virus genomes, and suggest that fission and fusion of virus genome segments or genes do not incur prohibitive adaptation costs.

We also detected multiple cases of structural gene module displacement with non-homologous counterparts. For instance, whereas members of *Potyviridae*, *Benyviridae* and *Matonaviridae* encode 3 unrelated CPs and form helical filamentous, rod-shaped or enveloped virus particles, respectively, some of the novel lineages branching near potyviruses, benyviruses and matonaviruses encode single jelly roll (SJR) CPs, and are thus expected to have non-enveloped icosahedral virions. Given the basal position of these virus lineages, it is likely that SJR CPs were ancestral in all three virus groups. Notably, in *f.0226.base−Beny* group, several viruses (e.g., ND_172503 and ND_246367) encode both SJR and tobacco mosaic virus (TMV)-like CPs that can be predicted to form icosahedral and helical virions, respectively (Figure S5. Genome maps). Apparently, in these viruses, the second CP gene has been acquired, but the ancestral one was also retained. This is likely to be a case of exaptation (refunctionalization) of one of the CPs, as previously described for closteroviruses, where two paralogs of the major CP and two unrelated structural proteins gained additional functions (Dolja et al., 2006). Non-homologous CP genes were also identified in the basal lineage to *Togaviridae* (*f.0271.base−Toga* and *f.0273.base−Toga*) and in *Virgaviridae*. Instead of the icosahedral capsid-forming chymotrypsin-like CP unique to *Togaviridae*, *f.0271.base−Toga* and *f.0273.base−Toga* encode a TMV-like CP, which is predicted to form rod-shaped helical virions. Thus, the TMV-like CP apparently emerged in a common ancestor of the *Hepelivirales* and *Martellivirales* orders, which include both animal and plant viruses. Conversely, in several identified *Virgaviridae* contigs (ND_191857 and ND_019381), the TMV-like CP was apparently replaced by structural proteins of *Kitaviridae*, including the major structural protein SP24 and a glycoprotein (Kuchibhatla et al., 2014).

Another notable replacement occurred in the *f.0268.base−Toga* group, where the structural module typical of togaviruses including the genes for CP and class II fusion protein was replaced by a class I fusion protein and M protein of nidoviruses (e.g., in ND_164660; Fig.5). An unexpected replacement of a membrane fusion glycoprotein was also identified in the *Xinmoviridae* family, where the characteristic class III fusion protein was replaced by a class II fusion protein, while retaining the typical mononegaviral nucleocapsid protein, thereby exemplifying the first member of the order *Mononegavirales* with a class II fusogen.

**Figure 5.**
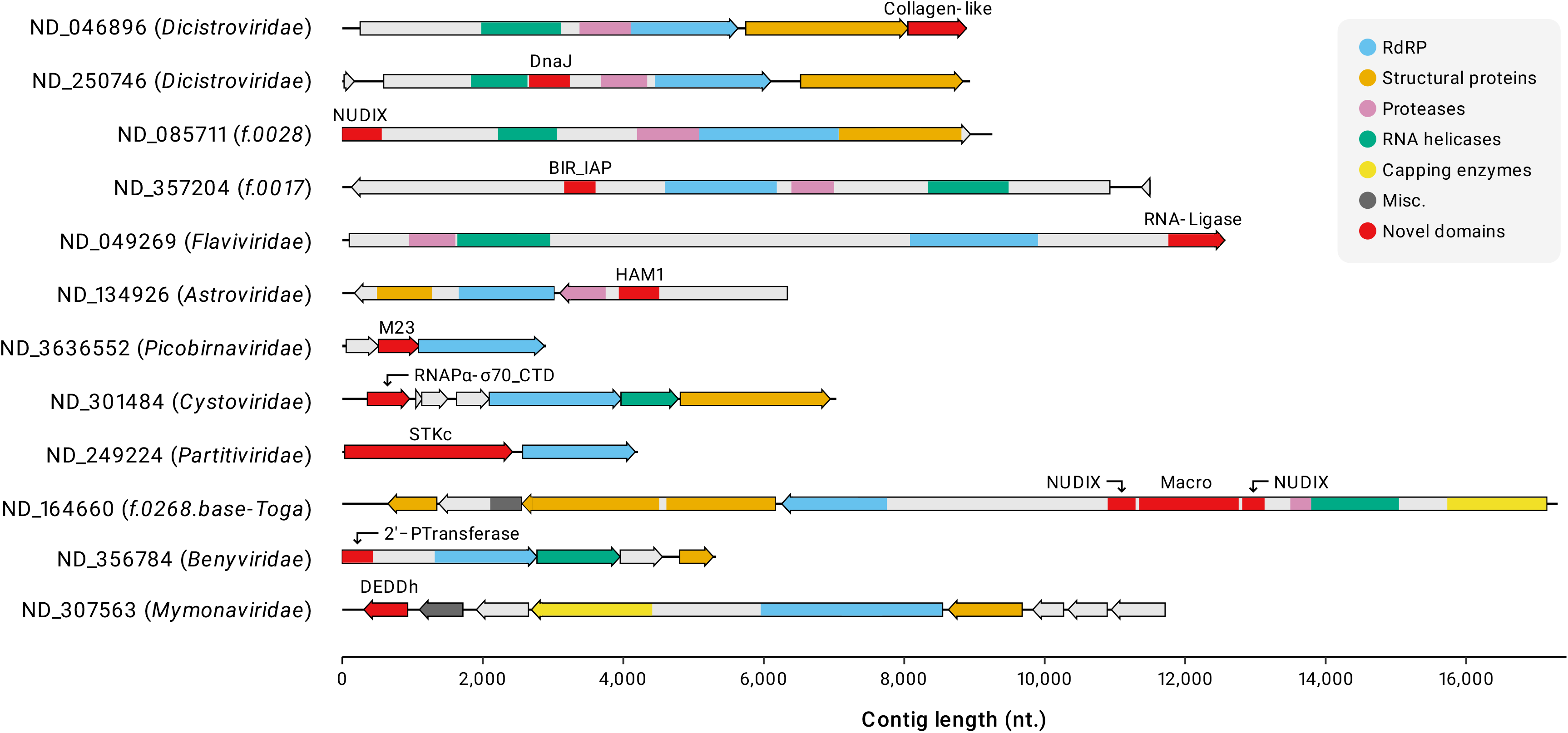
Diversity of new protein domains detected in RNA viruses. Homologous protein domains are shown as boxes of the same colours and the key is provided on the right side of the panel. Protein domains not commonly found in RNA viruses are shown in red and labelled above the corresponding boxes. Virus taxa and the unique contig identifiers are indicated on the left for each virus genome. The scale at the bottom shows the length (in nucleotides) of depicted genomes. Abbreviations: NUDIX, nucleoside diphosphate-X hydrolase; BIR_IAP, baculoviral IAP repeat (BIR) domains of the inhibitor of apoptosis (IAP); HAM1, inosine triphosphate pyrophosphatase; M15, M23 and M34, peptidoglycan digesting peptidases of MEROPS families M15, M23 and M34, respectively; RNAPα-σ70_CTD, a fusion of C-terminal domain of the α subunit of bacterial DNA-dependent RNA polymerase and C-terminal domain of σ70 factor; STKc, serine/threonine protein kinase; 2’−PTransferase, tRNA 2’-phosphotransferase; DEDDh, DEDDh-superfamily 3’-5’ exonuclease; RdRP, RNA-dependent RNA pol polymerase; Misc, miscellaneous.

We identified several virus groups basal to *Hypoviridae*, a family of capsid-less fungal viruses which do not have an extracellular infection stage. The newly discovered viruses encode CPs homologous to those of flexible helical viruses (*f.0066.base−Hypo*) or SJR CPs of icosahedral viruses (*f.0067.base−Hypo*, *f.0068*.*base−Hypo*, *f.0069.base−Hypo*), pinpointing the likely capsid-encoding ancestors of this virus family. Similarly, we identified SJR CP-encoding members of the family *Deltaflexiviridae* (e.g., ND_196199 and ND_246366 from *f.0215.base−Deltaflexi*), another family of capsid-less fungal viruses. The CPs of these new deltaflexiviruses were most similar to those of tymoviruses, which together with the filamentous alpha-, beta- and gammaflexivuses, and capsid-less deltaflexiviruses comprise the order *Tymovirales*, suggesting that *Deltaflexiviridae* evolved from a member of *Tymoviridae*, likely following the switch to fungal hosts.

Overall, our findings highlight the modular nature of RNA virus genomes, where genome replication and virion formation modules follow distinct evolutionary trajectories and are readily exchanged between distantly related viruses with radically different virions. Furthermore, the recurrent appearance of the SJR CP in base lineages of several groups of structurally diverse viruses is in agreement with the proposed origin of most eukaryotic RNA viruses from a simple ancestor that most likely encoded RdRP and SJR CP (Koonin et al., 2020; Wolf et al., 2018).

### Motif swapping in RdRPs

A notable feature of the RdRPs is swapping of the catalytic motifs (less accurately denoted “domain permutation”) that has been originally detected in the families *Permutotetraviridae* and *Birnaviridae*. The subsequent metagenomic analysis showed that the RdRP motifs were swapped on multiple occasions, in particular, in the large branch within *Kitrinoviricota* denoted the Yangshan assemblage (Wolf et al., 2020). In this work we identified 2,241 permuted RCR90 sequences and show that motif swapping was ancestral in two distinct class level clades (Fig. 2). One is a candidate class *c.0017* in *Pisuviricota* that consists of 264 RCR90 sequences and includes *Permutotetraviridae* and *Birnaviridae*, as well as 14 other family-level clades (*f.0088-f.0101*). The other is class *c.0032* in *Kitrinoviricota* consisting of 1,676 RCR90 sequences and covering 8 families (*f.0167-f.0174*). This class is part of a large branch of *Kitrinoviricota*, including classes *c.0031-c.0037*, to which the majority of the Yangshan assemblage genomes belong.

Additionally, in two smaller families within the Yangshan assemblage, *f.0158* and *f.0161*, the majority of the RdRPs (10 out of 12 and 89 out of 110, respectively) contained swapped motifs; in the latter family, swapping either occurred on two independent occasions or one ancestral swap reverted to the canonical configuration multiple times. Additionally, 23 families, 17 of these within *Kitrinoviricota*, included small clades RdRPs with swapped motifs, bringing the total number of independent swaps to at least 27 (25 independent events at or below family level plus permutations ancestral to two classes). This list includes a small clade of two permuted RCR90 sequences in *Botourmiaviridae* (*Lenarviricota*), which is so far the only RdRP motif swap outside of *Pisuviricota* and *Kitrinoviricota*.

### Expansion of the protein domain repertoire of RNA viruses

In addition to the RdRP, RNA viruses encode multiple other proteins and domains. Some, such as CPs, helicases and proteases of different families, are found in highly diverse viruses, whereas most are confined to relatively narrow virus groups (Koonin and Dolja, 1993; Wolf et al., 2020). Given the typical small genome size of RNA viruses, their protein domain catalogue expands relatively slowly with the increase of diversity measured by the RdRP analysis. In particular, in a previous metatranscriptomic study that doubled the global RNA virome, only three domains that were not previously identified in RNA viruses were detected (Wolf et al., 2020). Here, we performed an extensive search for protein domains encoded in the predicted RNA viral contigs. Briefly, we combined several protein domain databases (Pfam, COG, CDD, RNAvirDB2020, etc.) and ran multiple rounds of sensitive HMM profile searches (Söding, 2005), starting with the unmodified public profiles. This was followed by a multi-step custom procedure that involved profile curation, case by case examination of search results, and eventually, the generation of a new curated and annotated HMM profile database used for an expansive six-frame-translation HMMsearch (see Methods and Fig. S3).

The frequencies of the detected domains followed the expected power law-like distribution, where numerous domains were identified only in narrow groups of RNA viruses, and only a few widespread, hallmark domains, RdRP being by far the most common one, followed by multiple types of capsid proteins (CP_SJR, CP_levi), RNA helicases (SF1, SF2, SF3), and serine and cysteine proteases (Fig. S7). In addition to the aforementioned lysis-related domains, a number of new domains were detected that, to the best of our knowledge, have not been previously observed in RNA viruses.

In particular, we identified several domains, which are likely to modulate virus-host interactions and suppress the host antiviral response. Several members of the *Tobaniviridae* (order *Nidovirales*) which is known to primarily infect vertebrates, encode homologs (HHpred P=100%) (Zimmermann et al., 2018) of the cytokine receptor-associated Janus kinase (JAK) TYK2, which is activated upon receptor triggering and subsequently activates immune responses, including innate immunity against viral infection (Haan et al., 2006). Viruses evolved multiple mechanisms to disrupt the JAK-dependent signalling, emphasising the importance of this pathway in antiviral defence (Nan et al., 2017). The typical JAKs encompass FERM (four-point-one, ezrin, radixin and moesin), SH2 (Src homology 2), pseudokinase and kinase domains, where the pseudokinase auto-inhibits the kinase domain, thereby regulating the phosphorylation activity (Lupardus et al., 2014). The tobanivirus kinases apparently lack the FERM and SH2 domains and might function as dominant negative inhibitors of the cellular JAKs through their pseudokinase domains. Partitiviruses are the only other group of RNA viruses that encode a homolog (e.g. ND_249224 in Fig.5) of serine/threonine kinases, although the partitivirus protein is not specifically related to JAKs.

Members of the families *f.0059*.base-Poty and *f.0167* encode homologs of the cytokine receptors of the tumour necrosis factor receptor superfamily (TNFRSF), primarily involved in apoptosis and inflammation (Gravestein and Borst, 1998). The virus-encoded homologs could function as decoys of the cellular counterparts, sequestering the cytokines. Insect viruses of the family *Dicistroviridae* as well as several basal lineages of the *Solinviviridae* (*f.0024.base-Solinvi*, *f.0014.base-Solinvi*, *f.0017.base-Solinvi*, *f.0018.base-Solinvi*) and *Polycipiviridae* (*f.0008.base-Polycipi*) encode homologs of the BIR-domain (Baculovirus Inhibitor of apoptosis protein Repeat; Pfam: PF00653) proteins, first discovered in baculoviruses and subsequently identified in other, mostly, insect viruses with large dsDNA genomes (Lupardus et al., 2014). Notably, in the bona fide inhibitors of apoptosis (IAP), the BIR domain is followed by a RING ubiquitin ligase domain, whereas in the absence of the RING domain, BIR domain proteins function in cell cycle control (Clem, 2015; Gravestein and Borst, 1998). In dicistroviruses, the BIR domain is followed by a conserved cysteine-rich domain of unknown function (YP_009333511-212-260). The function of RNA virus BIR domains remains unclear and might be linked to regulation of cell death, cell cycle or both.

NUDIX superfamily hydrolases are widely represented in all three domains of life and in dsDNA viruses (Vasudevan and Ryoo, 2015). The virus homologs encoded by members of the *Poxviridae*, *Mimiviridae* and *Asfarviridae* are mRNA decapping enzymes that accelerate viral and host mRNA turnover, and promote the shutoff of host protein synthesis (Kago and Parrish, 2021; McLennan, 2006; Silke and Vaux, 2001). To our knowledge, no NUDIX hydrolases have been reported in RNA viruses to date. Instead, it has been demonstrated that human NUDIX hydrolase 2 (NUDT2) is involved in antiviral immunity by trimming 5′-phosphates from virus PPP-RNA, thereby handing the virus RNA off for degradation by the 5′-3′ exonuclease XRN1 (Parrish et al., 2007, 2009). We identified NUDIX hydrolases in viruses from 13 widely different families, including *Flaviviridae*, *Nodaviridae*, *Cystoviridae* and several new RNA virus families. Similar to the NUDIX hydrolases of dsDNA viruses, those encoded by RNA viruses of eukaryotes might be decapping enzymes, but in bacterial viruses, a different function is expected given that bacterial mRNAs lack eukaryotic-type caps.

One of the most widespread new domains identified in RNA viruses is the J domain so named after DnaJ (Hsp40), a co-chaperone that is conserved throughout the cellular tree of life as well as in some DNA viruses, and is important for protein translation, folding/unfolding, translocation and degradation, chiefly, through stimulating the ATPase activity Hsp70 chaperones (Kampinga et al., 2019; Laudenbach et al., 2021). Cellular DnaJ proteins play multiple roles during various stages of infection with diverse RNA and DNA viruses (Goodwin et al., 2011; Iyer et al., 2021; Taguwa et al., 2015). Notably, polyomaviruses, besides making extensive use of cellular DnaJ proteins (Goodwin et al., 2011), encompass a J domain at the N-termini of their replication proteins (large T antigens). This J domain is implicated in cell cycle control and cellular transformation (Brodsky and Pipas, 1998). We identified J domains in 11 diverse RNA virus families from the phyla *Kitrinoviricota* and *Pisuviricota,* which encode co-chaperones as part of their polyproteins. These RNA virus J domains could facilitate polyprotein folding and processing, and/or virion assembly. To our knowledge, involvement of DnaJ-like proteins in RNA virus replication has not been reported although closteroviruses encode a homolog of HSP70, the chaperone partner of DnaJ, which is involved in virion morphogenesis, itself incorporates into virions and facilitates virus intercellular transport (Alzhanova et al., 2007; Avisar et al., 2008).

We also identified several enzymatic domains that are likely to be involved in RNA repair and metabolism, including RtcB-like 3’-phosphate RNA ligase (Hughes et al., 2020); HAM1-like pyrophosphatase that hydrolyzes non-canonical purine nucleotides (Simone et al., 2013); DEDD-superfamily 3’-5’ exonuclease that could be involved in immune suppression, as demonstrated for arenaviruses (Hastie et al., 2011); and tRNA 2’-phosphotransferase implicated in tRNA splicing (Sawaya et al., 2005). In cellular organisms, the latter enzyme is often encoded together with NADAR (<u>NAD</u> and <u>A</u>DP-ribose) domain proteins implicated in NAD metabolism, in the context of RNA processing (de Souza and Aravind, 2012). NADAR domains have been originally detected in RNA viruses of the *Roniviridae* family and giant dsDNA viruses (de Souza and Aravind, 2012). We identified NADAR domains in RNA viruses from 12 families, emphasising the potential importance of this domain for RNA virus replication. An intriguing possibility seems to be that the RtcB-like RNA ligase and tRNA 2’-phosphotransferase repair tRNAs that might be cleaved in response to virus infection. Collagen is another domain that has not been previously detected in RNA viruses, but was identified in new members of the established families of *Dicistroviridae* (*Pisuviricota*), and *Tombusviridae* (*Kitrinoviricota*). In both families, a distant homolog of a “Collagen-like” protein family (KOG3544) was identified. While yet-unknown in RNA viruses, some giant dsDNA viruses, for example, mimiviruses, encode multiple collagen domain-containing proteins as well as enzymes involved in post-translational modification of collagen (Luther et al., 2011). The functions of viral collagens remain unknown.

Another unexpected finding is a protein encoded by bacterial *Cystoviridae* that contains an N-terminal domain related to the C-terminal domain (CTD) of bacterial sigma factors of the sigma70 family, a subunit of the cellular RNA polymerase holoenzyme that directs bacterial core RNA polymerase to specific promoter elements (Paget and Helmann, 2003). The CTD of this cystoviral protein corresponds to the C-terminal region of the bacterial RNA polymerase alpha subunit. The CTDs of sigma70 and RNA polymerase alpha are known to interact (Chen et al., 2003), suggesting that the viral protein reconstitutes this interaction interface and might participate in transcriptional takeover during infection, potentially overcoming the host antiviral defences.

Overall, as one could expect, the discovery of these diverse domains that have not been previously detected in RNA viruses and are restricted to one or several lineages implies multiple mechanisms of virus-host interaction, and in particular, counter-defence, that remain to be investigated experimentally.

### Alternative genetic codes in RNA viruses

Previous metatranscriptome analyses have shown that several groups of RNA viruses from different phyla utilise non-standard genetic codes suggesting that they infect hosts with matching codes, such as ciliates (Wolf et al., 2020). In the present analysis, of the 402,747 representative contigs, 25,080 (5.8%) showed evidence of alternative genetic codes, as attested by the presence of canonical STOP codons within the RdRP core domain region (see Methods). In most cases, it was impossible to reliably identify the specific alternative code, but where such an assignment was feasible, the most common codes were 6 (UAA and UAG stop codon recoded to Gln) and 14 (UAA and UGA recoded to Tyr and Trp, respectively, along with recoding of three sense codons), that have been identified in ciliates (Ring and Cavalcanti, 2008) and flatworm mitochondria (Ross et al., 2016), respectively. Many DNA phages appear to actively reprogram the host cell’s translation machinery for the phage protein synthesis via the use of alternative codes (Ivanova et al., 2014; Yutin et al., 2021). In contrast, non-standard genetic codes employed by RNA viruses are likely to represent adaptation to the host translation machinery as is the case for multiple isolated mitoviruses within *Lenarviricota* crown group that utilise the mitochondrial genetic code (UGA recoded from stop to Trp) and replicate inside the host mitochondria (Nibert, 2017). As expected, in this work, many virus lineages within the vast (5,006 RCR90 sequences) family *Mitoviridae* (*Lenarviricota*) were found to use code 4 that is common in fungal mitochondria. The largest of these lineages (399 RCR90 sequences) consisted entirely of viruses predicted to use this code including about 50 viruses from known fungal hosts. Overall, more than 50% of the viruses currently included in the family *Mitoviridae* use code 4.

Apart from the mitoviruses, contigs with alternative genetic codes were detected in most of the large RNA virus groups, typically, at frequencies of a few percent. To get a deeper insight into the spread and evolutionary conservation of alternative genetic codes in diverse virus groups, we identified virus lineages enriched in such codes at 50% or more throughout the phylogenetic tree of the RdRPs (Table S5, green arcs in Fig. 2). No phylogenetically coherent signal of alternative code usage in *Duplornaviricota* and *Negarnoviricota* was detected. In contrast, in *Pisuviricota*, 19 families, most of them new, typically contained one or two relatively small branches with 8-30 representatives associated with apparent protist codes (UAA, UAG or both reassigned to code for an amino acid). Among these, the *Dicistroviridae* family (∼2,000 RCR90 sequences altogether) stood out with 12 such branches, implying that some of the dicistroviruses previously thought to infect primarily arthropods may instead be viruses of protists, possibly, arthropod-associated ones. In *Kitrinoviricota*, a surprising, remarkable distribution of alternative codes prevalence was observed: 7 of the families in this phylum exhibited patterns similar to those in *Pisuviricota*, that is, one or several small branches with presumably protist codes used by the majority of the RCR90 sequences comprising the respective branch. In contrast, the remaining 7 families consisted exclusively (*f.0150, f.0177-f.181*) or primarily (*f.0176*) of viruses using alternative protist-like codes (about 900 sequences in total). In accordance with a recent report on several virus groups of this phylum (Yangshan assemblage) from a single brackish water habitat using protist-like genetic codes (Wolf et al., 2020), the present analysis of thousands of metatranscriptomes from diverse environments suggests that *Kitrinoviricota* include a substantial, previously unsuspected diversity of protist viruses.

The overwhelming majority of the samples analysed in this work originated from diverse microbial communities, but 109 RNA virus sequences were detected in transcriptomes of monocultures of microeukaryotes from the MMETSP project (Keeling et al., 2014). Several of these microeukaryotes utilise non-standard genetic codes. It is therefore plausible that viruses identified in these samples infect the respective microeukaryotes. Specifically, of the 57 MMETSP samples, 3 microeukaryotes were described as utilising non-standard genetic codes. In the respective transcriptomic assemblies, we identified two contigs of the same sequences from the family *f.0072.base-Flavi,* which were predicted to utilise genetic code 6 (ciliate), in line with the code used by the microeukaryote from the same samples (*Eukaryota; Sar; Alveolata; Ciliophora; Intramacronucleata; Spirotrichea; Oligotrichia; Strombidiidae; Strombidium*).

## Discussion

In recent years, metagenomes and metatranscriptomes have become the principal sources of DNA and RNA virus discovery, respectively (Call et al., 2021; Simmonds et al., 2017). Here we analysed more than 2.5 million RNA virus contigs recovered from 3,598 diverse metatranscriptomes. This analysis resulted in a 9-fold-increase in the number of 90% RvANI clusters (between the species and genus ranks), and an about 5-fold increase in the total phylogenetic depth, an almost 6-fold increase in the numbers of representative RdRP sequences (RCR90), and a 5-fold increase in the number of putative taxa at the levels from family to class. In contrast, at the phylum level, the RNA virus taxonomy remained essentially stable, with the exception of adding two new candidate phyla to the previously established five. Hence, across all criteria used to assess genome diversity, the present analysis yielded at least a 5-fold expansion of the global RNA virome.

Most of the previous assignments of RNA virus families to phyla remained stable through this extended analysis. However, notable exceptions were identified as well. Thus, the family *Cystoviridae* expanded by an order of magnitude and relocated from *Kitrinoviricota* to *Pisuviricota*, where it now forms a strongly supported clade with other dsRNA viruses, namely, picobirnaviruses and partitiviruses. Given the greater reliability of phylogenetic analysis with a substantially expanded family and considering the plausibility of the monophyly of these three groups of dsRNA viruses, the current position of *Cystoviridae* is likely to be valid and is expected to remain stable in future analyses. However, the placement of several other families, most notably, *Flaviviridae*, was shown to be unstable. Although this family also moved from *Kitrinoviricota* to *Pisuviricota*, in this case, the actual affiliation remains uncertain. Nevertheless, it is worth noting that the new affiliation of the flaviviruses with *Pisuviricota* is supported by their genome expression mode, namely, large polyprotein processing by a chymotrypsin-like protease, which is shared among *Picornavirales*, *Nidovirales* and *Flaviviridae*, as well as the presence of related superfamily II helicases in *Flaviviridae* and *Potyviridae* (Koonin and Dolja, 1993; Koonin et al., 2015).

Whereas the division of the kingdom *Orthornavirae* into phyla appears to be robust, the resolution of the phylogeny of the RdRP near the root might be insufficient to decipher the relationship among the phyla. The previously proposed scenario of the origin of dsRNA viruses from within positive-sense RNA viruses on multiple independent occasions and of negative-sense RNA viruses (*Negarnaviricota*) from the *Duplornaviricota* (Wolf et al., 2018) remains biologically plausible. However, phylogenetic analysis of the substantially expanded set of RdRPs reported here failed to vindicate this scenario in its entirety although multiple origins of dsRNA viruses were supported. The basal position of *Negarnaviricota* observed here is difficult to explain definitively. Given that negative-sense RNA viruses are primarily found in multicellular eukaryotes (Koonin et al., 2015), it is unlikely that this phylogenetic position of *Negarnaviricota,* albeit robust to the tests we performed, reflects the actual course of RNA virus evolution, and a yet unaccounted for artefact of deep phylogenetic analysis appears to be a more likely explanation. However, it is impossible to rule out that host assignments of newly identified members of *Negarnaviricota,* including those discovered here, change this perspective. In contrast, the basal position of *Lenarviricota* in the tree rooted with RT seems likely to reflect the actual origin of the rest of the RNA viruses from a common ancestor with this phylum within the bacterial domain. This scenario appears particularly plausible considering the major expansion of the bacterial RNA virome revealed here. Generally, considering the size and diversity of this dataset, it appears likely that the information contained in the sequences of the conserved regions of the RdRP is indeed insufficient to resolve the deepest relationships among RNA viruses. This problem will merit revisiting once sufficient diversity of RdRP structures accumulates, providing for a refined alignment and possibly a better phylogenetic resolution.

The analysis described here has a substantial impact on our understanding of the global diversity and ecology of RNA viruses. Most notably, the long-standing and perhaps enigmatic bias in the RNA virome towards eukaryote-infecting viruses (Koonin et al., 2015), is now effectively eliminated. Apart from the major expansion of the diversity of levi-like viruses, we obtained indications that multiple additional groups of viruses, in particular, picobirnaviruses and several clades of partitiviruses, likely infect bacteria. One of the key lines of evidence supporting this possibility is the discovery of numerous CRISPR spacers targeting RNA viruses, both members of *Leviviricetes* and a new group of putative RNA phages within partitiviruses. Prior to this work, there was little to no evidence of RNA virus targeting by CRISPR although targeting of RNA transcripts of DNA viruses by type III and type VI CRISPR-Cas systems as well as spacer acquisition from such transcripts mediated by the RT associated with the type III adaptation module is well documented (Abudayyeh et al., 2016; Mohanraju et al., 2016; Silas et al., 2017). Even with the findings of this work, the coverage of known RNA phages with CRISPR spacers remains low which is likely to reflect the strict requirement for RT for CRISPR adaptation to RNA viruses, and possibly the high evolution rate of RNA viruses which may prevent the identification of near-exact matches between contemporaneous RNA phages and CRISPR spacers. Notwithstanding the major expansion of the diversity of viruses that are known or strongly suspected to infect prokaryotes, the majority of metatranscriptomes from various environments remain dominated by viruses assigned to eukaryotes. Thus, although the prokaryotic RNA virosphere appears to be far more diverse than previously suspected and now covers a substantial fraction of the overall RNA virus diversity, quantitatively, RNA viruses remain a relatively small component of the prokaryotic virome in most biomes, while being much more prevalent in the eukaryotic virome. A corollary is that analysis of metatranscriptomes from specialised ecosystems might yield many unusual groups of RNA viruses infecting prokaryotes as exemplified by the putative new phyla p.0002 and RvANI90_0011770 described here.

The present results strongly suggest that drastic host shifts, known as horizontal virus transfer (HVT), between distantly related hosts, even crossing the prokaryote-eukaryote divide, is a major route of RNA virus evolution, as previously proposed for viruses infecting eukaryotes (Dolja and Koonin, 2018). The HVT events likely occurred on multiple, independent occasions within different phyla, classes and possibly, even orders of RNA viruses. In that regard, the small group of viruses, for which multiple CRISPR spacer matches were detected and that therefore was tentatively assigned to the *Roseiflexus* bacterium as the host, is notable. This narrow virus group from a unique habitat, roughly constituting a genus, is lodged deeply within partitiviruses, many of which are known to infect fungi, plants and invertebrates (Cross et al., 2020; Shi et al., 2016; Vainio et al., 2018). Conceivably, many more such groups of RNA viruses infecting prokaryotes and pointing to possible HVT events remain to be discovered in diverse, isolated, still poorly sampled habitats.

The robust phyletic relation between *Leviviricetes* and the eukaryotic viruses in the *Mitoviridae*, *Narnaviridae*, and *Botourmiaviridae* was unequivocally supported in this work, and these eukaryotic virus families were substantially expanded, especially *Botourmiaviridae*, which became the second-largest family within the kingdom *Orthornavira*. These findings support the notion that bacterial viruses directly seeded a major component of the eukaryotic virome, possibly, via the mitochondrial endosymbiont (Koonin et al., 2020; Wolf et al., 2018). Endosymbiosis could be the biological mechanism underlying other, perhaps numerous HVT events as well. Indeed, endosymbiotic bacteria that are common in protists, fungi, animals (in particular, insects) and plants (McCutcheon and Moran, 2011; Moran and Bennett, 2014) might be habitual vehicles of HVT. In the case of *Lenarviricota*, the direction of HVT from bacteria to eukaryotes appears indisputable, but on other occasions, capture of eukaryotic viruses by bacteria cannot be ruled out, especially, taking into account the cases of apparent gene transfer from protist-infecting viruses to bacteria (Lurie-Weinberger et al., 2010). Mechanistically, HVT between bacteria and eukaryotes might present a challenge given the substantial differences in translation mechanisms. The evolutionary scenario for *Lenarviricota* illustrates how this challenge can be overcome by loss and recapture of genes encoding capsid proteins. Furthermore, interdomain HVT is likely to be facilitated for viruses with genomes consisting of multiple monocistronic RNA segments as in the case of partitiviruses.

In addition to the major expansion of the global RNA virome, this work also substantially expands the catalogue of protein domains encoded in RNA virus genomes including several domains with predicted activities that have not been previously observed. The common theme among these domains that are each represented in narrow lineages of RNA viruses appears to be counter-defence via diverse molecular mechanisms. These findings seem to indicate that, despite their typically smaller genomes, RNA viruses are more similar to DNA viruses with respect to exaptation of host genes than previously appreciated.

In summary, the collection of RNA virus sequences reported here far exceeds in diversity those described previously, suggests many new functionalities for RNA virus proteins, and can be expected to become a major resource for RNA virus research. The present results greatly expand the kingdom *Orthornavira*, while introducing relatively minor changes into the latest taxonomic scheme, thus supporting its general robustness. While this article was in the final stages of preparation, a massive metatranscriptome analysis reporting numerous novel RNA viruses has been published (Edgar et al., 2022). A thorough comparison of the results of the two studies is a major task in itself that remains for future analyses, and will hopefully take us one step closer to a comprehensive exploration of the Earth RNA virome.

## Methods

### Metatranscriptome acquisition

The identification of RNA viruses was performed on a total of 5,150 publicly available metatranscriptomes that were retrieved from IMG/M in January 2020 (Chen et al., 2021; Mukherjee et al., 2021) (Table S7 - sample metadata). Sequences shorter than 1,000 nt. or encoding rRNA genes were discarded and the remaining contigs were dereplicated at 99% sequence identity.

### Primary Filtering process

For convenience, we summarised the final tools and cutoffs of the Primary and secondary filtration process in Table S6 - threshold summary.

To filter out sequences that were highly unlikely to represent RNA viruses, we compared the obtained metatranscriptome contigs to a compendium of DNA sequences built from 1,831 metagenomes originated from the same studies as 1,306 of the metatranscriptomes. We selected metagenomes that shared the metadata attribute of “Study_ID” with the 5,510 metatranscriptomes in the GOLD portal (Mukherjee et al., 2021) as these DNA assemblies would cover a similar range of habitats as the analysed metatranscriptomes. Using multiple sequence search tools (specifically, MMSeqs2 (nucleic - nucleic (search type 3) (Hauser et al., 2016; Steinegger and Söding, 2017), DIAMOND (translated nucleotide versus the IMG sourced DNA metagenomic predicted ORFs (diamond blastx) (Buchfink et al., 2015), and NCBI BLAST (nucleic - nucleic - blastn) (Altschul et al., 1997; Camacho et al., 2009; Johnson et al., 2008)) in an iterative manner, we identified and excluded metatranscriptomic contigs that matched sequences in the DNA sequence dataset (Figure 1A), based on the assumption that RNA viruses would not be present in DNA assemblies — which would be comprised of cellular organisms, DNA-based mobile elements, and integrated retroviruses. The iterative search was performed such that each iteration gradually increased the search sensitivity (e.g., through decreased word length (BLASTn) and higher sensitivity value (MMSeqs2 “--sensitivity”)), while discarding all sequences from the metatranscriptomes collection that produced reliable matches to sequences in the “DNAome”, before advancing with the filtered output to the next iteration.

### Secondary Filtering process

To further filter the contig set, we supplemented the above filtering process output with 5,954 RNA viral sequences from reference databases and performed an additional iterative filtering procedure using public databases (NCBI NT/NR and IMG/VR) as the DNA set. To prevent the exclusion of *bona* fide RNA virus sequences, we masked entries of the public databases that matched reference RNA viruses from subsequent iterations. All discarded contigs were aggregated and supplemented with manually identified DNA encoded contigs, creating a database of “false positives”, that was used to further filter the metatranscriptome dataset through exclusion of sequences with producing passable matches to the “false positive” set. The procedure of collecting the discarded matches to further refine the working set was repeated three times.

### Estimation of DNA remnants in intermediate sets

To evaluate remnants of DNA sequences in the working set through the filtration process, we routinely analysed random contig subsets by (1) computing the RdRP to reverse-transcriptase domain ratio as a proxy to the RNA virus to DNA-encoded contigs; (2) manually inspecting the presence of the most frequent non-RNA virus-related domains. Of note, several specific domains recurred frequently during this performance evaluation, and manual examination revealed these to be domains of known repeats. Mostly, these contigs were fully populated with matches to such repeat domains, and that these had cellular matches in the public DBs, whose alignment values were just below our reporting or acceptance criteria. Hence, we decided to discard these contigs if they were completely coding for multiple repeats, as there would be no sufficient coding space for these to encode an identifiable RdRP.

Following the below RdRP identification step (described in the section below) approximately 130 reverse-transcriptases had passed the various filtration processes and were manually removed. MMSeqs2, the PFamA Database (Mistry et al., 2021) and the RdRP and RT collection from Wolf et al 2018, were used in all the profile searches performed in this evaluation.

### RdRP identification

Previously published multiple sequence alignments of RdRPs and reverse-transcriptases (Wolf et al., 2018 and Wolf et al., 2020) were formatted as tool-specific subject databases, and employed as queries to search a sequence database consisting of the 6-frame end-to-end translations of contigs passing the above-described filtering processes, using PSI-BLAST, hmmsearch, DIAMOND and MMSeqs2. To estimate the desired search cutoffs, we supplemented the query set with non-RdRP sequences likely to produce false matches (termed “true-negative” set), constructed as follows: (1) using a large set of RdRPs as queries for an hhsearch (from the HH-Suite) against the PDB70 database (2019), collecting all matches of bitscore ≥ 20 that were not from RNA viruses, that aligned with at least 2 RdRPs; (2) fetching PDB entries clustered with those at 70% identity, (via ftp://resources.rcsb.org/sequence/clusters/bc-70.out); (3) fetching Pfam entries relating the resulting PDB IDs, and sequences linked to the Pfam entries; (4) collapsing highly similar sequences to a single representative (MMSeqs2 minimum coverage: 100%, minimum ident.: 90%). Subject RdRP profiles capable of producing alignments to any sequence from the “true-negative” set were discarded. Otherwise, acceptance criteria for the RdRP profiles searches were: profile coverage ≥ 50%, E-value ≤ 1e-10 and score ≥ 70. These stringent parameters were then fine tuned to represent the best possible value a non-RdRP sequence was able to achieve.

Subsequently, reliable RdRP matches were trimmed to the approximate core domain, which we operationally defined as motif A–D (see “Motif A–D identification” below). The extracted RdRP core sequences were pre-clustered (CD-HIT, coverage ≥ 75%, % ID ≥ 90) (Fu et al., 2012), passed to an all vs. all (DIAMOND BLASTp) run, formatted for use with MCL using mcxload (--stream-mirror --stream-neg-log10 -stream-tf “ceil(200)”), clustered (MCL, Inflation value between 3.6 and 2.8), aligned (MUSCLE), and formatted as profile databases as described above (Altschul et al., 1997; Buchfink et al., 2015; Edgar, 2021; Enright et al., 2002; Steinegger and Söding, 2017). This process was repeated twice. Subsequently, contigs with putative RdRPs were used to recover additional contigs from the entire metatranscriptomic collection, which were highly similar yet shorter than the initial search length criteria (see below “Comprehensive identification” for details). Of the resulting collection, sequences covering ≥ 75% of an RdRP profile, or with identifiable motifs A–D, were considered sufficiently complete for downstream phylogenetic analysis.

### Identification of the RdRP catalytic motifs A**–**D

A custom motif library (available in the project Zenodo archive, see code and data availability) was built by semi-manual partitioning of the previously published RdRP MSAs. To identify the motifs along the individual RdRP sequences, a similar iterative search as described above for the full length RdRP domain was performed.

### Correction of putative frame shifts

Several thousand of the contigs identified contained a clear RdRP domain signature on more than one frame, commonly separated by a single nucleotide. In order to avoid the omission of these signatures as simple incomplete, we addressed these in two manners: (1) if any of one of the signatures covered ≥75% of the subject RdRP profile, or coding for the desired catalytic motifs A–C, that signature would be used; or (2) by concatenation of the two signatures into a single amino acid sequence.

### Contig set augmentation with published genomes

To assess the novelty of our findings in terms of the number and diversity of newly predicted viral genomes, and in order to avoid the exclusion of established viral lineages that may be underrepresented in environmental metatranscriptomes, we aggregated and compiled a collection of “previously published’’ viral genomes termed “Reference Set”. These include RdRP-carrying sequences identified in NCBI’s NT database (NCBI Resource Coordinators, 2018), as well as sequences not indexed (at the time of writing) in such public databases, that were identified in several previous large scale and notable RNA virus surveys and transcriptomic atlases. Our criteria for addition of these supplementary sequences required that they originate from peer reviewed publications, and that all underlying sequences were entirely publicly available, with no restrictions. The NCBI NT sequences were identified via an RdRP scan procedure similar to the procedure described above (see RdRP identification). The previously published set was made from an expansive set of Leviviricetes described by Callanan et al (Callanan et al., 2020), the “Yangshan-assemblage” and other described by Wolf et al (Wolf et al., 2020), and the proposed Plastroviruses group described by Lauber et al (Lauber et al., 2019), as well as several RdRPs identified in the ocean atlas of genes (Salazar et al., 2019; Tara Oceans Coordinators et al., 2018). Following their aggregation, these sequences underwent a similar procedure described for the metatranscriptomic sequences identified in this work (i.e. length filtration, clustering, and RdRP core domain extraction). The eventual sequence set was labelled as “Known” (i.e. not novel), and noted as such in the data generated by this work (e.g. branch colour in Fig. 1). The processed “supplemental sequence set” was merged into the main sequence set (those identified in this study) and the combined set (termed “VR1507”) was used in all downstream analysis (phylogenetic reconstruction, domain analysis etc).

### Comprehensive identification of RNA virus contigs across metatranscriptomes

Because metatranscriptome assemblies can often yield incomplete genomes that would not fulfil the criteria for *de novo* RdRP detection (see above), we used the “VR1507” contig set (see above), we initiated a secondary “sweeping” scan for additional RNA virus contigs from the non-clustered, non-filtered (length, DNA similarity, RdRP presence) “bulk-set” of metatranscriptomic contigs (Fig. 1). To this end, the “VR1507” was used as bait for highly similar contigs in the “bulk-set”, using an non-sensitive mmseqs search (mmseqs search --search-type 3 --min-aln-len 120 --min-seq-id 0.66 -s 1 -c 0.85 --cov-mode 1) followed by stringently filtering the recovered matches (E-value < 1e-9, Identity > 95%, Target-Coverage ≥ 95%). These criteria were selected as a quality assurance measure, so that the recovered contigs would be mostly contained within the “VR1507” contig counterpart (this large expansive data set is available in project’s Zenodo repository see code and data availability). This envelopment criteria was added to avoid capture of chimeric, or otherwise uncertain, nucleic regions. The filtered bulk contig set was combined with “VR1507”, and consisted of 2,658,344 contigs (termed “Add1507”).

### Phylogenetic reconstruction

We selected a diverse set of representative RdRPs for the phylogenetic analysis by performing a preliminary MMSeqs2 clustering run (see Table S8 - Clustering information), on a subset of the sequences which contained complete or near-complete RdRPs. These representatives were termed RCR90, and went through several iterations of clustering (MMSeqs2 with sequence identity threshold of 0.5), alignment (MUSCLE5) (Edgar, 2021)and profile-profile comparison (HHSEARCH) (Steinegger et al., 2019), as described below. “Permuted” RdRPs (sequences with transposed motif C, following the C-A-B-D configuration) were identified and “de-permuted” (i.e. the loop, containing motif C, was cut from the sequence and reinserted downstream from the motif B). Once all identified sequences with transposed motifs were brought into the canonical A-B-C-D configuration, the following procedure was employed to produce a multiple sequence alignment consisting of all RCR90 set:

- Sequences were clustered using MMSeqs2 with sequence identity threshold of 0.3; sequences in the resulting 4,514 clusters were aligned using MUSCLE5; profile-profile comparison of the cluster alignments using HHSEARCH produced a 4,514×4,514 distance matrix (the distances were estimated as *d*_AB_ = -ln(*S*_AB_/min(*S*_AA_, *S*_BB_)), where *S*_AB_ is the HHSEARCH score for comparison of the profiles A And B); a maximum-linkage tree was produced from the distance matrix using the R function hclust();
- The tree was cut at the depth threshold of 1.5, producing 1,360 subtrees;
- Each of the subtrees was used as a guide to hierarchical alignment of the corresponding profiles using HHALIGN, producing 1,360 alignments;
- 1,360 consensus sequences (excluding sites with more than 2/3 of gap characters) were extracted from these alignments and aligned using MUSCLE5;
- Each position in the alignment of consensus sequences was expanded to the corresponding column of the original alignment, producing an alignment of 77,510 RdRps (where the original RdRp sequences were reduced to a set of positions, matching their local consensus);
- Sites with >90% of gap characters were removed from this alignment; the resulting alignment was aligned with the alignment of ten RTs (five group II intron sequences and five non-LTR retrotransposon sequences) using HHALIGN.

The alignment of RdRps and RTs was used to reconstruct an approximate maximum likelihood tree using the FastTree (V.2.1.4 SSE3, Price et al., 2010) program (WAG evolutionary model, gamma-distributed site rates) and rooted between RTs and RdRps.

### Taxonomic affiliation of clades

Tree leaves with existing taxonomic information were identified by mapping (MEGA-BLAST, E-value < 1e-30, query coverage ≥ 95%, subject coverage ≥ 95%, Alignment length > 200, Identity ≥ 98%, (Alignment_length)/Query_length > 0.95) VR1507 sequence set to the latest ICTV data at the time of analysis (July 20, 2021 release of the Virus Metadata Repository (VMR) file, corresponding to MSL36, and available at https://talk.ictvonline.org/taxonomy/vmr/m/vmr-file-repository/13175). Overall, 2,765 contigs were mapped, and the ICTV taxonomic information was cloned to the VR1507 queries based on the highest score. For the reminder of VR1507 contigs, we performed a similar procedure using the NCBI’s NR database (these amount to an additional 6,878 mapped contigs, though a non-negligible amount of those lacked taxonomic information or matched abolished taxonyms). The procedure to establish the taxonomic affiliation of internal nodes on the tree (i.e. clades) relies on the above taxonomic assignment of reference tree leaves, as well all on two principles:

- All sequences, descending from the last common ancestor of reference leaves, assigned to a taxon *T*, also belong to taxon *T*; sequences descending from deeper tree nodes, do not belong to taxon *T* and, therefore, and should be assigned to a new taxon (taxa) of the same rank;
- The depth, at which a tree clade splits into taxa of the given rank, is defined by existing taxa of the same rank and is locality-dependent (e.g. the characteristic depths of families could be different for different phyla);

Application of these principles assumes that the existing taxonomy is non-contradictory with respect to the tree, i.e. the reference sequences, assigned to taxa, form monophyletic clades that are non-overlapping and non-nested within the same rank (e.g. a family clade can’t be embedded into another family). An inspection of the taxonomic affiliation of reference leaves showed that this assumption, while typically satisfied, is violated in multiple places. This necessitates disentangling the conflicting relationships first. To this end, the following procedure was applied to all taxa of the given rank (i.e. separately for phyla, classes, etc):

- The tree was pruned to contain only leaves with this rank defined (e.g. all leaves without a family assignment are stripped); leaf weights (*w_i_*) were derived from the pruned tree;
- For each taxon *T*, present in the tree, the total weight of leaves in this taxon was calculated (W_T_ = Σ*w_i_* across the leaves, assigned to *T*);
- For any tree clade in the tree, the total weight of leaves in this clade was calculated (W_C_ = Σ*w_i_* across the leaves, belonging to *C*);
- For each combination of clade *C* and taxon *T*, the clade-taxon weight was calculated (*W_CT_* = Σ*w_i_* across the leaves, belonging to *C* and assigned to *T*); then a precision-like and recall-like measures can be calculated (*P_CT_* = *W_CT_* / *W_C_* and *R_CT_* = *W_CT_* / *W_T_*) and combined into a quality index *Q_CT_* = *P_CT_* * *R_CT_*.
- For each taxon *T*, present in the tree, the clade *C_T_* = argmax *Q_CT_* was identified as the “native” location of the taxon *T* (the clade, where the maximum weight of taxon *T* is concentrated with the minimal intrusion of other taxa); leaves, belonging to clade *C_T_*, but not assigned to *T*, and leaves, assigned to *T*, but not belonging to clade *C_T_*, were labelled as “intruding” or “outlying” respectively;

All tree-incompatible taxonomic assignments were examined and resolved. In most cases the most agnostic way to resolve the conflict was used (i.e. stripping the taxonomic labels from the corresponding leaves). In one case, most of the families within *Timlovirales* order of *Lenarviricota*, were found to be nested inside a very deep-branching family of *Blumeviridae*. For the purpose of this work, we retained the *Blumeviridae* label on the largest clade of *Timlovirales* that didn’t have conflicting family assignments and removed the *Blumeviridae* label from the rest of *Timlovirales*. In a few other cases where small families were wholly nested into larger ones (e.g. a solo leaf classified as *Sunviridae* inside a large *Paramyxoviridae* clade) the embedded family label was removed for the purpose of subsequent analysis and restored post hoc. Once the taxonomic labels of all leaves were brought into compatibility with the tree, the following procedure was performed to assign new taxonomic labels to unlabeled leaves for each taxonomic rank separately:

- All nodes of the tree were assigned depth, defined as the longest node-to-leaf path across all leaves, descending from this node;
- In the full tree of 77,510 leaves the last common ancestor node of each taxon was determined; depths of the taxa, defined as the depth of the LCA node plus the length of the incoming tree edge, was recorded; all unlabelled leaves, descending from the taxon LCA, were assigned to this taxon;
- All clades outside of existing taxa were isolated; for each such clade the depths of all existing sister taxa were determined; if a clade has only one sister taxon, the search for the closest relatives was extended toward the root until at least another related taxon was identified; the threshold depth was calculated as the average for the set of related taxa;
- Clades outside of existing taxa were dissected at the threshold depth; each resulting (sub)clade was assigned to a new taxon of the given rank;
- New taxa that have a single existing taxon as a sister are labelled as associated with this taxon.

The novel taxa were given names, indicating rank (i.e. prefixed by *p*, *c*, *o, f* and *g* for phylum, class, order, family and genus respectively), followed by an ordinal number for new taxa of this rank, and optionally, terminated with a label for taxa that are associated with a previously described taxon (e.g. *f.0127.base-Noda* is the 127th new family that is basal to *Nodaviridae* in the RdRP tree).

### Robustness of deep phylogeny

To assess the robustness of deep phylogenetic reconstruction, the following procedure was performed:

- a list of 201 families with at least 20 RCR90 sequences was collected
- a random representative of each family and from RT set was sampled
- a sub-alignment of 202 sequences for the sample was extracted from the master alignment
- a phylogenetic tree was reconstructed using the IQ-Tree program (Nguyen et al., 2015) with an automatically selected best fitting model
- 100 independent samples were analysed in the following manner:

First, clades with the highest quality index (QI, described above in the Taxonomic affiliation of clades section) were identified for each of the five known phyla; the quality index values were used as a measure of the phylum monophyly under the subsampling. Families, involved in breaking the monophyly of the respective phyla (note that a leaf can be both an outlier with respect to its own phylum and an intruder into another phylum), were recorded.

Second, the subsampled trees were collapsed to the phylum level; 15 (out of 100) trees with paraphyletic phyla were excluded (those, where e.g. the highest-quality clade for Pisuviricota was embedded within the highest-quality clade for Kitrinoviricota). An extended majority-rule consensus tree was constructed for the remaining 85 trees with (largely) monophyletic phyla using the IQ-Tree program; branch support values were multiplied by 0.85 (the fraction of such trees among the whole sample).

### Assignation of individual contigs to RCR90 clusters

Once the novel areas of the RCR90 megatree described above were fully populated by the major taxonomic ranks (Phylum→Genus), we proceeded to affiliate contigs from the larger VR1507 set (see above - contig sets). Contig affiliation was performed in a gradual manner by separation into the following 4 levels:

Level A. are contigs encoding the RdRPs used to create the tree. Level B. consists of contigs encoding RdRPs with exceptionally high amino acid identity to RdRPs from level A, (via best BLASTp match with Identity ≥90%, Query-Coverage ≥75%, and E-value < 1e-3). Level.C consisted of contigs from the same RvANI90 cluster (see definition below) as contigs from levels {A, B}, and Level D. consists of contigs sharing high nucleic similarity to those from levels {A - C}, (via best dc-MEGABLAST hit at Identity ≥90%, Query-Coverage ≥75% OR Nident ≥ 900nt and E-value < 1e-3). Based on the distribution of ICTV-labelled RdRPS in the above noted levels, we estimate that the majority contigs affiliated in this manner, would roughly share the same taxonomic ranks down to Genus level.

Of note, for level C., we devised custom measurement unit, RvANI, which is an extension of standard average nucleic identity (ANI) clustering, designed to accommodate the fragmented nature of metatranscriptomic assemblies, thus avoiding an overestimation of novelty caused by the relatively low pairwise coverage of related sequences. Briefly, RvANI is calculated as follows: Initially, mmseqs is used to calculate all pairwise sequence alignments in the contig set, which are then used for the traditional ANI and alignment fraction (AF) calculations, where:

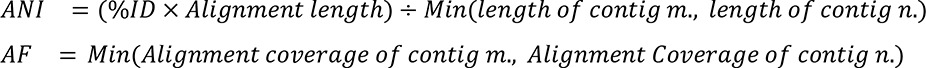

Given all pairs of ANI and AF (for prokaryotes 95-96% ANI is the commonly accepted species boundary, with similarly granular definitions for certain viruses (Nayfach et al., 2021; Richter and Rosselló-Móra, 2009) clusters are defined as connected components in a nucleic similarity graph pruned for pairwise alignments with ANI ≥90% and AF ≥90%. RvANI corrects for uneven genome coverage in metaTs by reinserting specific pairwise alignments to the pruned nucleic similarity graph, even if their AF is below the required cutoff, as long as the underlying pairwise alignment fulfil these criteria: %ID ≥ 99,

Alignment Length ≥ 150[bp], and the alignment occurs between the edge of the contigs, i.e. the alignment covers the 5’ or 3’ termini of each contig. See Table S8).

### Identification of reliable CRISPR spacer hits

RNA virus sequences were compared to predicted bacteria and archaea CRISPR spacer sequences to (i) identify which viruses may infect a prokaryotic host, and (ii) possibly predict a specific host taxon for these viruses. First, non-redundant RNA virus sequences were compared to 1,568,535 CRISPR spacers predicted from whole genomes of bacteria and archaea in the IMG database (Chen et al., 2021) using blastn v2.9.0 with options “-dust no -word_size 7”. To minimise the number of false-positive hits due to low-complexity and/or repeat sequences, CRISPR spacers were excluded from this analysis if (i) they were encoded in a predicted CRISPR array including 2 spacers or less, (ii) they were ≤ 20bp, or (iii) they included a low-complexity or repeat sequence as detected by dustmasker (v1.0.0) (Morgulis et al., 2006) (options “-window 20 -level 10”) or a direct repeat of ≥ 4bp detected with etandem (v6.6.0.0) (Rice et al., 2000) (options “-minrepeat 4 -maxrepeat 15 -threshold 2”). To link RNA viruses to CRISPR spacers, only blastn hits with 0 or 1 mismatch over the whole spacer length were considered. The spacer and array with hits were further inspected to check (i) whether the spacers were of consistent length throughout the array, and (ii) whether Cas and/or RT genes were found in the putative host genome, and if so whether these were adjacent to the CRISPR array with the hit. To expand the search for CRISPR link beyond bacteria and archaea for which a draft genome is available, we next used the same approach to compare non-redundant RNA virus sequences to 53,372,161 CRISPR spacers predicted from metagenome assemblies available in the IMG database. Spurious spacers were filtered out using the same methods as for the genome-derived CRISPR arrays (see above), and only hits for which the RNA virus and the CRISPR spacers originated from the same ecosystem (as defined in the GOLD database) were retained. Since CRISPR spacer arrays are often assembled on short contigs without any other gene, we used the repeat sequence of the arrays to link them to a putative host. Repeat sequences from metagenome-derived CRISPR arrays with at least 1 hit to an RNA virus sequence were compared to all IMG Bacteria and Archaea genomes using blastn (v2.9.0) with options “-perc_identity 90 -dust no -word_size 7”. The location of these hits in the putative host genome was then checked for the presence of a predicted CRISPR spacer array, Cas genes, and RT genes. When individual RNA virus sequences or spacers were putatively linked to multiple host genomes, these were prioritised based on the following criteria: (i) the spacer array is identified next to an RT-encoding CRISPR array, (ii) an RT-encoding CRISPR array is identified elsewhere in the genome, (iii) the spacer array is identified next to a Type III CRISPR array, (iv) a Type III CRISPR array is identified elsewhere in the genome, (v) another type of CRISPR array is identified in the genome, and (vi) no identifiable Cas gene can be identified in the genome.

The spacer content of CRISPR arrays encoded by *Roseiflexus sp. RS-1* in Mushroom Spring was further studied as follows. First, the CRISPR arrays of 17 metagenomes sampled from microbial mats in Mushroom Spring (Table S3) were specifically assembled using the dedicated tool Crass v1.0.1 with default parameters (Skennerton et al., 2013). Next, all arrays based on repeats corresponding to known CRISPR arrays in *Roseiflexus sp. RS-1* (Table S3) were identified and the corresponding spacers collected and filtered as previously described. RNA virus sequences as well as DNA virus sequences from the IMG/VR v3 database (Roux et al., 2020) were compared to this database of *Roseiflexus sp. RS-1* spacer arrays using blastn (v2.9.0) with options “-dust no -word_size 7”. Sequences from putative RNA phages infecting *Roseiflexus sp. RS-1* were first identified based on hits to ≥ 1 RS-1 spacer with ≤ 1 mismatch across the whole spacer length. For these selected phages, hits with up to 4 mismatches across the spacer length were then collected to enable the detection of more distant virus-spacer hits.

Candidate capsid segments of *Roseiflexus sp. RS-1* clade *genPartiti.0019 viruses* were identified based on 3 criteria: spacer match to the RNA-targeting CRISPR array, no corresponding DNA sequence, and high coverage correlation to ≥ 1 RdRP contig across the metatranscriptome time series. First, a similar blastn comparison to Crass-assembled spacers (blastn with options “-dust no -word_size 7” and ≤ 1 mismatch allowed) was used to identify putative capsid-encoding contigs i.e., excluding all contigs encoding an RdRP or a CRISPR array, in the same metatranscriptomes targeted by the *Roseiflexus sp. RS-1* Type III-RT CRISPR array (n=3,958). Next, candidates with ≥ 1 spacer match were compared to all contigs from Mushroom Spring DNA metagenomes (blastn (v2.9.0) with options “-task megablast -max_target_seqs 500 -perc_identity 90”), and all candidates with a matching DNA contig (≥ 90% identity) were considered to be likely DNA phages and excluded (n=3,650). Finally, the coverage of all *genPartiti.0019* RdRP contigs and all candidate capsid segments was obtained using read mapping as described below (bbmap.sh (v.38.90) with options “vslow minid=0 indelfilter=2 inslenfilter=3 dellenfilter=3”), and candidates with a Pearson correlation of ≥ 0.9 across the 42 Mushroom Spring metatranscriptomes were retained as likely capsid segments (n=88). To evaluate the gene content of these capsid segments, cds were predicted de novo using Prodigal (v2.6.3) (Hyatt et al., 2010) (option “-p meta”), and clustered using a standard blast-mcl pipeline (blastp (v2.9.0) with default options, hits selected based on score ≥ 50, MCL clustering (v.14-137) with an inflation value of 2). For the three largest protein clusters, a sequence alignment was built using MAFFT v7.407, (Katoh and Standley, 2013) and used as input to an hhsearch against the virus-focused uniprot public database (uniprot_sprot_vir70), and a custom database made from capsids of known partitiviruses and picobirnaviruses (available in the project’s Zenodo repository, see code and data availability “Partiti_Picob_CP.tar.gz” and PC1_PROMALS3D_new.hhr).

### Habitat distribution and relative abundance estimation

For visualisation purposes, location, ecological, and taxonomic information for each metatranscriptome were obtained from the IMG and GOLD databases. Specifically, GPS coordinates and ecosystem classification were obtained from GOLD, with the ecosystem information further grouped in custom categories (Table S7). To roughly estimate the host diversity present in each metatranscriptome, the taxonomic information of all contigs as predicted by the IMG annotation pipeline (Clum et al., 2021) was queried at the domain level, i.e. Bacteria, Archaea, Eukarya, and Viruses. The ratio between the number of contigs assigned to Bacteria and Archaea and the number of contigs assigned to Eukarya was then used as a proxy to determine “Prokaryote-dominated” from “Eukaryote-dominated” datasets. Specifically, datasets with a ratio of Eukaryote-affiliated to Prokaryote-affiliated contigs ≤ 0.3 or ≥0.7 were considered as “Prokaryote-dominated” or “Eukaryote-dominated”, respectively, while other datasets were considered as “Mixed”. The map was drawn using the packages matplotlib v3.3.4 and basemap v1.2.2 for python 3.8.5 (Hunter, 2007).

For read mapping, a dereplicated set of RNA virus sequences (95% ANI over 95% AF, established using CheckV anicalc.py and aniclust.py scripts (Roux et al., 2020)), was established, hereafter “NR-mapping” dataset. Quality-trimmed reads (sensu (Clum et al., 2021) from 3,998 metatranscriptomes (Table S7) were then mapped to this dataset as follows. First, contigs from each metatranscriptome were compared to the NR-mapping dataset using blastn v2.9.0+ (E-value ≤ 0.01). All contigs with cumulated blast hits of ≥ 90% average nucleotide identity covering ≥ 80% of the shortest sequence were considered as putative RNA viruses. All reads mapping to contigs identified as putative RNA viruses and all unmapped reads were extracted from the existing IMG read mapping information, and mapped de-novo on the NR-mapping dataset using bbmap v38.81 (Bushnell, 2014) with the following options: “vslow minid=0 indelfilter=2 inslenfilter=3 dellenfilter=3”. This step was done to reduce the computing time and the risk of false-positive mapping by excluding all reads mapping to non-viral metatranscriptome contigs. The resulting bam files were then filtered with FilterBam (https://github.com/nextgenusfs/augustus/tree/master/auxprogs/filterBam) retaining only mapping at ≥50% identity and ≥50% coverage, and genomecov from bedtools v2.30.0 (Quinlan, 2014) was used to calculated the average coverage depth for each contig in each sample. The relative proportion of a taxon was then calculated as the cumulated coverage for the taxon members divided by the total accumulated coverage of all predicted RNA virus contigs in this dataset.

### Genetic code assignment and ORF calling

Presently, ORF identification software designed for diverse metagenomic data are limited to the standard genetic code (11) or the Mold mitochondrial genetic code (4) (opted when the predicted ORFs are unnaturally short). To identify clades likely to use alternative genetic codes, we extracted the RdRp core footprints and scanned them for in-frame standard stop codons.

We first separated all RdRP-encoding contigs into two subsets: “standard” and “non-standard” if any canonical stop codons occurred within the narrow coordinates of the RdRP core. Then, the “standard” set was subjected to metaprodial CDS prediction using default parameters (via Prodigal’s (v2.6.3) metagenomic mode (“anonymous”)) (Hyatt et al., 2010). In the “non-standard” subset, the stop codon usage patterns were aggregated across the contigs, associated with each tree leaf, and classified into “mitochondrial” (using UGA as a sense codon), and “protist” (other patterns). Prevalence of patterns (relative frequency among the descendant leaves) was calculated for internal tree nodes; clades with high prevalence were noted and investigated. For practical purposes, the ORFs predictions of the “non-standard” subset were performed by using the first genetic code enabling the entire RdRP core to be translated. Cases for which none of the available genetic codes enabled the uninterrupted translation of the RdRP core were assigned the general “non-standard” value, and were predicted using the mitochondrial genetic code (4).

To discard the possibility of active recoding of tRNAs by these predicted RNA viruses, the VR1507 set was subjected to a single pass of tRNAscanME2 (Chan et al., 2021), using the “global” flag (for non-specific domain of life tRNA prediction). No tRNAs were identified on any of the viral contigs predicted to use an alternative genetic code, suggesting these are most likely an adaptation to their host rather than an element of a virus-host arms race, as seen in some dsDNA phages (Ivanova et al., 2014).

### RBS identification and quantification

Using VR1507 as input, the RBS quantification was performed as described in Schulz el al. 2020. Briefly, Prodigal (v2.6.3) was run as described above (see “Genetic code assignment”) (Hyatt et al., 2010; Schulz et al., 2020), we then sourced the “rbs_motif” field from Prodigal’s GFF output files, and classified the different 5’ UTR sequences as either “SD” (for motifs similar to AGGAGG, the canonical Shine-Dalgarno), “None” and “Other” (for details, see code and data availability, “RBS_Motif2Type.tsv”). Then, for each contig, we defined the “%SD” as the ratio between all “SD” ORFs, and all ORFs with a true start (i.e. not truncated by the contigs’ edge, field “start_type” different from “Edge”).

### Domain annotation

To perform an initial domain annotation of the proteins encoded by RdRP-containing contigs, we used hmmsearch (from the HMMER V3.3.2 suite) (Finn et al., 2011; Wheeler and Eddy, 2013) to match these proteins to HMMs gathered from multiple protein profile databases (PFam 34, COG 2020 release, CDD v.3.19, CATH/Gene3D v4.3, RNAVirDB2020, ECOD 2020.07.17 release, SCOPe v.1.75) (Andreeva et al., 2014, 2020; Cheng et al., 2015; Galperin et al., 2021; Lu et al., 2020; Mistry et al., 2021; Sillitoe et al., 2021; Wolf et al., 2020). We supplemented this set of HMMs with a custom collection of profiles with bacteriolytic functions (termed “LysDB” - available in the project’s Zenodo repository, see code and data availability). LysDB was built from (1) manually reviewed profile entries from public databases which we could link to GO terms related to cell lysis by viruses, or virus exit from host cell, and (2) custom profiles for “Sgl” proteins, which were experimentally demonstrated by Chamakura *et. al* to induce cell lysis (Chamakura et al., 2020). Additionally, we used InterProScan (v.5.52-86.0) to scan the protein sequences using MobiDBLite (v2.0), Phobius (v.1.01), PRINTS (v. 42.0), TMHMM (v.2.0c) (Attwood et al., 2012; Jones et al., 2014; Käll et al., 2004; Kall et al., 2007; Krogh et al., 2001; Potenza et al., 2015).

Because the public protein profile databases that were used for initial annotation might contain HMMs that represent polyproteins, which span multiple functional domains, we developed and employed a procedure to identify such profiles which were masked from the subsequent annotation process. For this procedure, we first used the hmmemit command to convert HMMER profiles into multiple sequence alignments, which were then used as input to an all-versus-all profile comparison performed using HH-Suite. Next, putative polyprotein profiles were identified by flagging the profiles that encompassed at least two other non-overlapping profiles (“get_polyproteins.ipynb” script, see data and code availability). The unmatched regions between the polyprotein domains were extracted to create a set of conserved, yet unknown domains, termed “InterDomains”. Additionally, profiles with over 1000 match states (defined as columns with less than 50% gaps) were manually examined using HHpred. Several of the identified polyprotein profiles were split into their constituent domains. Subsequently, all hmmsearch results were aggregated and profile matches were prioritised based on their classification level (uncurated profiles, or ones of unknown function (e.g. “DUF”) were deprioritized) and by their relative alignment statistics. To improve the quality of the functional annotation of the domain profiles and to assign functions to unannotated profiles we identified clusters of similar profiles (clans, hereafter). First, profiles with at least one hit in the initial (E-value ≤ 0.001) were extracted from their original DB, reformatted as HH-Suite’s HHMs (as described above) and used for an additional all-versus-all step. The output of this profile comparison was then used as input to a graph-based clustering process using the Leiden algorithm (“get_clan_membership.ipynb” script, see data and code availability), which identifies clans as communities of highly similar domains. Clan membership was then used to improve the coverage of the functional annotation by transferring annotation from functionally annotated profiles to other clan members. Briefly, this procedure followed a consensus-based label assignment. For example, a clan with 12 profiles labelled as “RdRP”, and 2 “unclassified” profiles, was set as an “RdRP’’ clan and the 2 unclassified members were reclassified as “RdRP”. Cases of conflicts were either left unresolved, or by opting to the lowest denominator. For example, a clan with 4 “unclassified” profiles, that also had 12 member profiles labelled “Super family 2 Helicase” and an additional 10 member profiles labelled “Super family 1 Helicases’’, was set to “Helicase-uncertain”, and this label was extended to those 4 “unclassified” members.

All subsequent profile matches passing a predefined cutoff (E-value ≤ e-7, score ≥ 9, alignment length ≥ 8[AA]). were used to generate a new custom profile database, in a process similar to the one used for RdRPs (see above). Only clusters with ≥ 10 sequences, sharing the same functional classification, were used to generate HMMs. This profile set was then supplemented by most of the profiles from the above-mentioned RNAVirDB2020 database, as well as several dozen select profiles from the other databases (this final profile database termed “NVPC” is available via the projects Zenodo repository, see data and code availability). Finally, we queried the six-frame translations of the **330k** contig set using hmmsearch as described above, using the new profile database. (Fig. S3 - Annotation pipeline). Subsequently, we generated tentative genome-maps for ≈4-20 representative contigs for each of the 400+ identified families (novel and established) using GGGenomes (https://github.com/thackl/gggenomes), which were then manually examined to identify novel domains as well as uncommon domain fusion and segmentations.

## Supporting information

Supplementary_archive

## Data and code availability

Custom analysis code used in this work is available under the open-source MIT Licence at https://github.com/UriNeri/RVMT. All data produced in this work is freely and fully available through several venues. In hope of providing a long lasting community resource, we have created an interactive web portal at https://riboviria.org that allows users to download portions of the data generated in this work based on phylogeny and data type (e.g., a subset of the domain annotations for all contigs affiliated with a certain family). Both programmatic and graphical access to the data are supported through the web portal. The website’s code is also available under the MIT Licence at https://github.com/Benjamin-Lee/riboviria.org. For all taxonomic levels, this platform includes raw nucleic sequence, phylogenetic trees, metadata, and annotations. To ensure data longevity, we have also permanently archived all the data produced in the course of this project in CERN’s Zenodo repository. The project as a whole (including future versions) is accessible at DOI:10.5281/zenodo.6091357 and the version used in the present article is accessible at DOI:10.5281/zenodo.6091357. This project is intended to serve as a community wide resource. As such, the Zenodo repository includes the additional information and various intermediary results and secondary analyses, such the predicted coding sequences, host assignments, phylogeny and taxonomic affiliation, raw domain HMM searches, additional domain profile databases generated in this work (e.g. alignments, HMMs, original seed sequences and predicted function) as well the nucleic sequences for both the expanded (2.6M metatranscriptome derived) contig set and the manually consolidated “Reference Set” (see Methods).

## The RNA Virus in metatranscriptomes consortium

Adrienne B. Narrowe, Alexander J. Probst, Alexander Sczyrba, Annegret Kohler, Armand Séguin, Ashley Shade, Barbara J. Campbell, Björn D. Lindahl, Brandi Kiel Reese, Breanna M. Roque, Chris DeRito, Colin Averill, Daniel Cullen, David A. C. Beck, David A. Walsh, David M. Ward, Dongying Wu, Emiley Eloe-Fadrosh, Eoin L. Brodie, Erica B. Young, Erik A. Lilleskov, Federico J. Castillo, Francis M. Martin, Gary R. LeCleir, Graeme T. Attwood, Hinsby Cadillo-Quiroz, Holly M. Simon, Ian Hewson, Igor V. Grigoriev, James M. Tiedje, Janet K. Jansson, Janey Lee, Jean S. VanderGheynst, Jeff Dangl, Jeff S. Bowman, Jeffrey L. Blanchard, Jennifer L. Bowen, Jiangbing Xu, Jillian F. Banfield, Jody W Deming, Joel E. Kostka, John M. Gladden, Josephine Z Rapp, Joshua Sharpe, Katherine D. McMahon, Kathleen K. Treseder, Kay D. Bidle, Kelly C. Wrighton, Kimberlee Thamatrakoln, Klaus Nusslein, Laura K. Meredith, Lucia Ramirez, Marc Buee, Marcel Huntemann, Marina G. Kalyuzhnaya, Mark P Waldrop, Matthew B Sullivan, Matthew O. Schrenk, Matthias Hess, Michael A. Vega, Michelle A. O’Malley, Monica Medina, Naomi E. Gilbert, Nathalie Delherbe, Olivia U. Mason, Paul Dijkstra, Peter F. Chuckran, Petr Baldrian, Philippe Constant, Ramunas Stepanauskas, Rebecca A. Daly, Regina Lamendella, Robert J Gruninger, Robert M. McKay, Samuel Hylander, Sarah L. Lebeis, Sarah P Esser, Silvia G. Acinas, Steven S. Wilhelm, Steven W. Singer, Susannah S. Tringe, Tanja Woyke, TBK Reddy, Terrence H. Bell, Thomas Mock, Tim McAllister, Vera Thiel, Vincent J. Denef, Wen-Tso Liu, Willm Martens-Habbena, Xiao-Jun Allen Liu, Zachary S. Cooper, Zhong Wang.

## Acknowledgments

The authors would like to thank Shai Zilberzwige-Tal, David Burstein, Adi Stern, Leah Reshef, Linda Yu and Omry Lieber for helpful discussions and support of this project.

U.G. and U.N. are supported by the European Research Council (ERC-AdG 787514). U.N. is partially supported by a fellowship from the Edmond J. Safra Center for Bioinformatics at Tel-Aviv University. Y.I.W. and E.V.K. are supported through the Intramural Research Program of the US National Institutes of Health (National Library of Medicine). V.V.D. was partially supported by NIH/NLM/NCBI Visiting Scientist Fellowship. The work of the U.S. Department of Energy Joint Genome Institute (S.R., A.P.C., I.M.C., N.I., D.P.E., N.K. and all JGI co-authors), a DOE Office of Science User Facility, is supported by the Office of Science of the U.S. Department of Energy under contract no. DE-AC02-05CH11231. M.K. was supported by l’Agence Nationale de la Recherche grants ANR-20-CE20-009-02 and ANR-21-CE11-0001-01. D.K. was funded by the European Social Fund under no. 09.3.3-LMT-K-712-14-0027. D. A. B. is supported by grant NNX16SJ62G from the NASA Exobiology program, and by grant DE-FG02-94ER20137 from the Photosynthetic Systems Program, Division of Chemical Sciences, Geosciences, and Biosciences (CSGB), Office of Basic Energy Sciences of the U. S. Department of Energy.

We gratefully acknowledge the contributions of many scientists and principal investigators, who sent extracted genetic material for isolate genomes, environmental metagenomes and metatranscriptomes, or sequencing results as part of the Department of Energy Joint Genome Institute Community Science Program, and allowed us to include in our study the RNA virus sequences detected in these publicly available data sets regardless of publication status.

