## Supplementary_archive for "A five-fold expansion of the global RNA virome reveals multiple new clades of RNA bacteriophages": Fig. S2 - Robustness estimates for the established Phyla.pdf

**A. Monophyly of phyla in 100 trees, reconstructed from subsampled alignments**

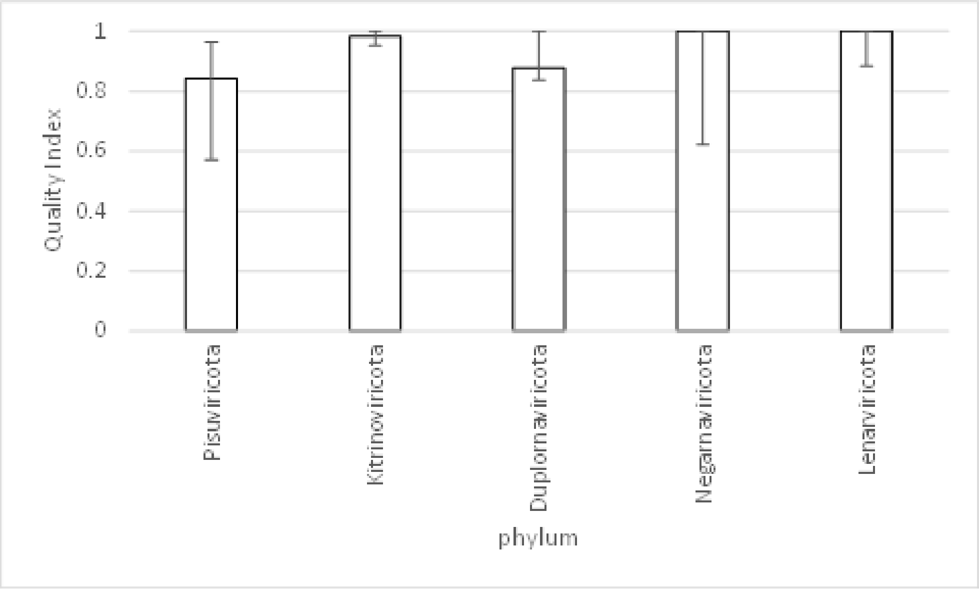


**B. Virus families most frequently involved in violations of phyla monophyly**

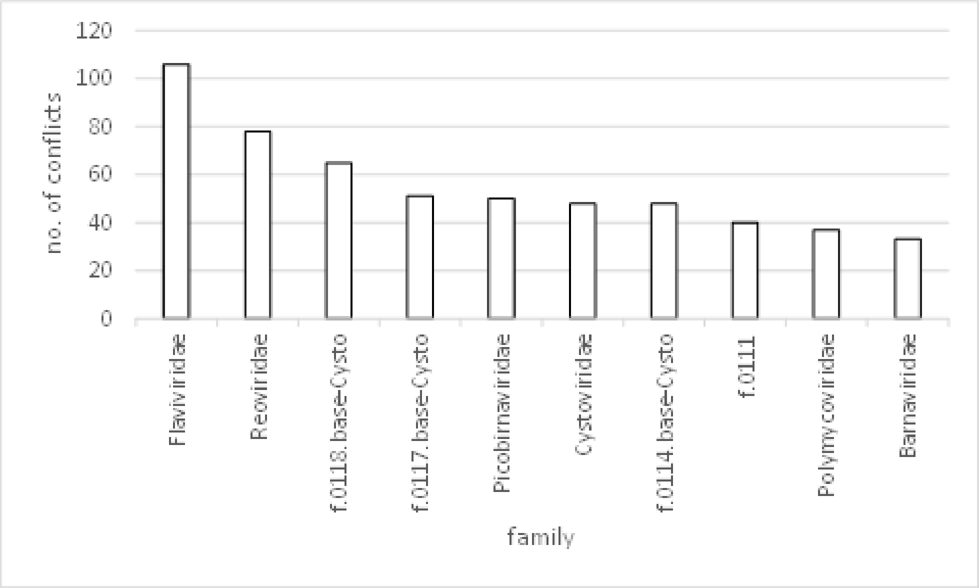


**C. Extended majority rule consensus tree for subsampled alignments**

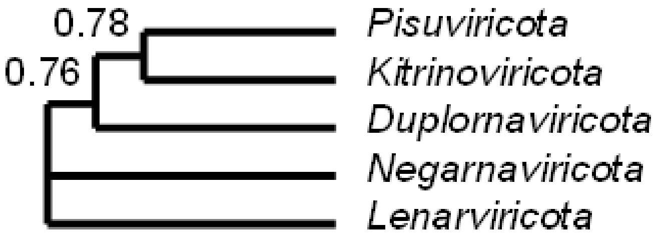
