## Supplementary_archive for "A five-fold expansion of the global RNA virome reveals multiple new clades of RNA bacteriophages": Supplementary figure legends.docx

### Supplementary information - Supplementary figure legends

**Figure S1:** Distribution of contigs in RCR90/RvANI clusters.

Each panel displays the total number of clusters (left panel RCR90, right panel RvANI90) on the horizontal axis (logarithmic scale) against their size (total number of memebring contigs) on the vertical axis (logarithmic scale).

**Figure S2:** Robustness of deep phylogenetic reconstructions.

A. Quality index (the product of the fraction of phylum members that form a monophyletic clade and the fraction of other phyla members in this clade). The bar shows the median value across 100 independent samples of one member of a family with at least 20 members; the whiskers indicate the 5% and 95% percentiles.

B. The virus families, most often involved in monophyly violations (where a leaf is either outside of the clade of its phylum or inside a clade of the other phylum). The number of violations is shown.

C. The extended majority consensus tree of the five previously known phyla. The consensus tree was recovered from 85 (out of 100) samples that have non-embedded monophyletic phyla and the support values were multiplied by 0.85.

**Figure S3:** *Roseiflexus* sp. RS-1 CRISPR arrays and related viruses. A. Map of the 4 CRISPR-Cas regions in *Roseiflexus* sp. RS-1 (NC_009523.1) including predicted CRISPR arrays (red diamonds) and Cas genes (colored genes). B. Coverage heatmaps across Mushroom Spring and Octopus Spring metagenomes, for spacers associated with *Roseiflexus* sp. RS-1 (see Table S3). Spacers matching predicted RNA phages are displayed on the left, and spacers matching DNA phages are displayed on the right for reference. 3. Example of alignment obtained with hhpred for a putative capsid protein from a predicted novel RNA phage infecting *Roseiflexus* sp. RS-1 and the closest publicly available homolog: Fig cryptic virus capsid protein.

**Figure S4:** Identification of different metatranscriptome types and associated virus types. A. Distribution of the ratio of viruses predicted to infect prokaryotic hosts across individual samples. B. Distribution of non-viral contigs affiliated as eukaryotes or prokaryotes (hosts) across samples, separated based on the protocol used to generate the metatranscriptome. The protocol information was obtained from the Gold database, and summarised as follows: “PolyA selection”: transcript enrichment based on polyA tail, “rRNA depletion”: use of a kit(s) and/or protocol(s) for depletion of rRNA templates, “Total RNA”: cDNA library prepared from the extracted RNA with no polyA selection or rRNA depletion step, “Unknown”: no information available. C. Relationship between the ratio of eukaryote/prokaryote RNA viruses (x-axis) and the ratio of eukaryote/prokaryote host contigs (y-axis). Each dataset type is presented in a separate panel.

**Figure S5:** Acquisitions and replacements of structural modules in RNA viruses.

“*Picobirnaviridae*; ND_299612” and “f.0226.base-Beny; ND_172503” exemplify fusions of genomic segments encoding capsid proteins (CP) and RdRPs, which are encoded on separate segments in previously described picobirnaviruses and benyviruses. “f.0066.base-Hypo; ND_049849” and “*Deltaflexiviridae*; ND_196199” encode TMV-like and single jelly roll (SJR) CPs, respectively, although other members of the respective families comprise capsid-less viruses. “*f.0271.base-Toga*; ND_366069” and “*Virgaviridae*; ND_191857” represent genomes with non-homologous replacements of the CP genes. In “*Xinmoviridae*; ND_221687”, class I fusion glycoprotein gene, typical of xinmoviruses and other members of the *Mononegavirales*, has been replaced by a gene encoding a class II fusion glycoprotein (CIIF). Abbreviations: Env, envelope protein; GP, glycoprotein; PRO-Pap/vOTU, papain-like protease; SF1, superfamily 1; Cap_MTase-GTase, capping enzyme with methyltransferase-guanylyltransferase activities.

**Figure S6:** Extended Annotation Pipeline.

Flowchart diagram visualising the procedures used in the domain identification and functional annotation sections of the project.

**Figure S7:** Identified Domain distribution.

The predicted viral function or structure of the final domain hits (vertical axis, slanted text labels), against the total number of reliable observed HMM search matches (horizontal axis, logarithmic scale).
