## Supplementary_archive for "A five-fold expansion of the global RNA virome reveals multiple new clades of RNA bacteriophages": Supplementary table legends.docx

### Supplementary information - Table legends

**Table S1:** RCR90 cluster data - Size, taxonomic lineage, genetic code, and permutations

Sheet "RCR90" columns:

1. RCR90 ID (cluster identifier)
2. "Native" contig ID (the original nucleic sequence identifier coding for the leaf)
3. Other contig IDs, associated with this tree leaf (comma-delimited list)
4. Other RdRp IDs, associated with this tree leaf (comma-delimited list)
5. Phylum
6. Class
7. Order
8. Family
9. Virus name (if exist)
10. Alt code - Genetic code information (empty, "Mito" or "Protist", with asterisk if it belongs to an alt-code clade).
11. Permuted - (empty, or "Permuted", with asterisk if belongs to a permuted clade)

Sheet "Families" columns:

1. Family name
2. Number of RCR90 clusters of this family

**Table S2:** Genome features of RNA viruses suggesting a prokaryotic host. The first tab (“RBS motifs”) indicates for each taxon with a putative prokaryotic host the total number of ORFs with a predicted Shine-Dalgarno-like motif, the total number of ORFs considered (ORFs predicted on the edges of contigs were excluded from this analysis), and the corresponding ratio of ORFs with predicted Shine-Dalgarno-like motifs. RBS motifs were predicted with prodigal, and the following motifs were considered as Shine-Dalgarno-like: 3Base_5BMM, 4Base_6BMM, AGG, AGGA, AGGA_GGAG_GAGG, AGGAG, AGGAG_GGAGG, AGGAG(G)_GGAGG, AGGAGG, AGxAG, AGxAGG_AGGxGG, GAG, GAGG, GAGGA, GGA, GGA_GAG_AGG, GGAG, GGAG_GAGG, GGAGG, GGAGGA, GGxGG. The second tab (“Bacteriolytic domains”) lists all predicted ORFs functionally annotated as encoding a lysis protein, for RNA virus taxa predicted to infect a prokaryotic host. For elevated e-values (> 0.01), the alignment was visually inspected to ensure that the active sites residues were conserved. The third tab (“CRISPR spacer hits”) lists all significant hits (0 or 1 mismatch) identified between RNA viruses with predicted prokaryotic hosts and the IMG spacer database. For each hit, the taxon and type of the corresponding CRISPR array (identified based on exact similarity of the repeat sequence) is indicated. “RT”: CRISPR array encoding a reverse-transcriptase.

Sheet “RBS signal” columns:

1. Taxon
2. #SD carrying ORFs
3. Total # of complete ORFs
4. %SD

Sheet “Bacteriolytic domain” columns:

1. ND
2. ORF
3. Taxonomic affiliation
4. E-value
5. Category
6. Domain hit

Sheet “CRISPR hits” columns:

1. RNA virus contig
2. CRISPR array type (NA: unknown)
3. Host taxon (NA: unknown, hit to unaffiliated metagenome-derived CRISPR spacer only)

**Table S3:** Detailed information linking new clades of partiti-like RNA viruses to *Roseiflexus* sp. RS-1 host. The first tab (“Full RdRP - CRISPR matches”) lists all hits (0 or 1 mismatches) identified between selected RNA viruses and CRISPR spacers associated with *Roseiflexus* sp. RS-1 arrays, obtained with the CRASS assembler. The second tab (“Capsid segment search”) lists the contigs identified as potential capsid segments based on (i) hits (0 or 1 mismatches) to the RT-encoding CRISPR array of *Roseiflexus* sp. RS-1 (column “Number of Spacer matches in NC_009523.1_3781897_3786321_CAS-III-B”), and (ii) high correlation to one of the RdRP-containing segments (column “Relative abundance correlation to closest RdRP”). The third tab (“Metatranscriptomes and metagenomes”) includes the list of datasets used in this analysis.

Sheet “Full RdRP - CRISPR matches” columns:

1. ## Genome
2. Spacer
3. Spacer length
4. Number of mismatches
5. Repeat
6. Array type
7. Spacer list of samples

Sheet “Capsid segment search” columns:

1. Potential capsid segment contig
2. Length
3. GC
4. Topology
5. Number of Spacer matches in NC_009523.1_3781897_3786321_CAS-III-B
6. Number of Spacer matches in other Roseiflexus sp. RS-1 arrays
7. Closest
8. Relative abundance
9. Correlation to closest RdRP
10. GC of closest RdRP

Sheet “Metatranscriptomes and metagenomes” Information derived from GOLD (“Genomes OnLine Database”) attributes, please refer to GOLD website for additional information.

1. IMG Taxon ID - Metatranscriptomic/genomic assembly identifier in the IMG/M database.
2. Dataset name
3. Sample Collection Date
4. Dataset type

**Table S4:** Characteristics of metatranscriptomes used for ecological distribution analysis. For each metatranscriptome, the type of protocol used is indicated when available, along with the type of dataset based on the number of contigs affiliated to eukaryotic or prokaryotic taxa, and the sample geographic coordinates when available.

Distribution of the ratio of viruses predicted to infect prokaryotic hosts across individual samples. B. Distribution of non-viral contigs affiliated as eukaryotes or prokaryotes (hosts) across samples, separated based on the protocol used to generate the metatranscriptome.

**Table S5:** virus lineages enriched in alternative genetic codes

Sheet "Families" columns:

1. Family name
2. Number of RCR90 clusters of this family
3. Number of RCR90 clusters with potential alt. codes (i.e. with standard stop codons in the RdRp core domain)
4. Number of RCR90 clusters with potential alt. codes that form high-density clades (frequency of alt-code sequences 0.5 and above)

Sheet "RCR90" columns:

1. RCR90 ID (cluster identifier)
2. Phylum
3. Class
4. Order
5. Family
6. Alt code - Genetic code information (empty, "Mito" or "Protist", with asterisk if it belongs to an alt-code clade)

**Table S6:** Discovery pipeline search thresholds

Values derived from the most lenient of the alignment acceptance thresholds used for each search and step, i.e. prior searches or subset based tuning rounds may have superseded by the values listed below. N.B. to reduce computational load, all DNA filtrations searches were run until the first reliable match per query (mmseqs max-accept 1, BLASTn max_target_seqs 1, DIAMOND --max-target-seqs 1).

Table columns:

1. Process
2. Step
3. Identity - minimal identity [%] for matches to be accepted as representing reliable alignments.
4. Alignment length - minimal length (either in nt or aa) for search results to be accepted as representing reliable alignment.
5. E-value - maxima E-value for matches to be accepted as representing reliable alignments.

**Table S7:** List of metatranscriptomes used in this study. For each metatranscriptome, the summarised ecosystem classification used in this study is indicated, along with the JGI proposal DOI and publication information when available.

Table columns:

1. Taxon_oid - identifier for the metatranscriptomic assembly in the IMG/M database.
2. Ecosystem - semi manual classification of the environment type from which the genetic material was sourced.
3. N representatives - number of unique RvANI90 representative contigs identified in the sample.
4. Total RNA virus contigs - number of all RNA viral contigs identified in the sample.
5. Proposal DOI (for proposals with ≥ 1 RNA virus detected)
6. Publication - Citable source to be used if describing the identified contigs.

**Table S8:** Cluster information

Data regarding the clustering procedures and results. Column are self explanatory, and provide the parameters, size distribution, description of input and output sets used as well as the code/tool for the different runs.

**Table S9:** Information regarding the project’s Consortium co-authorships:

1. Name
2. Email
3. Affiliation
4. Funding Statement
