## Supplementary figures and images for "A five-fold expansion of the global RNA virome reveals multiple new clades of RNA bacteriophages"

### Fig. S1 - RCR90and RvANI90 cluster size dist.pdf

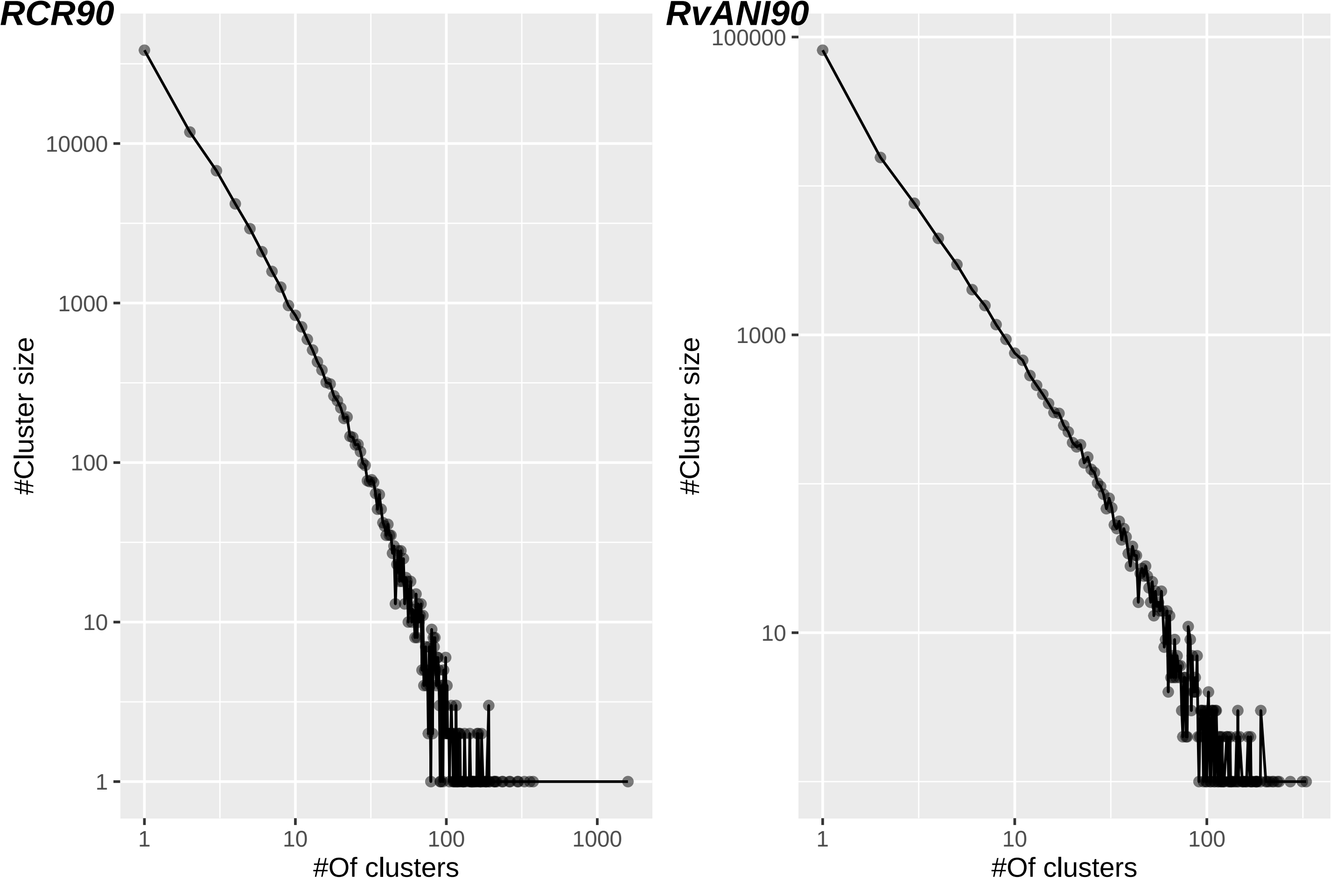

### Fig. S3 - DNA_CRISPR_heatmap.pdf

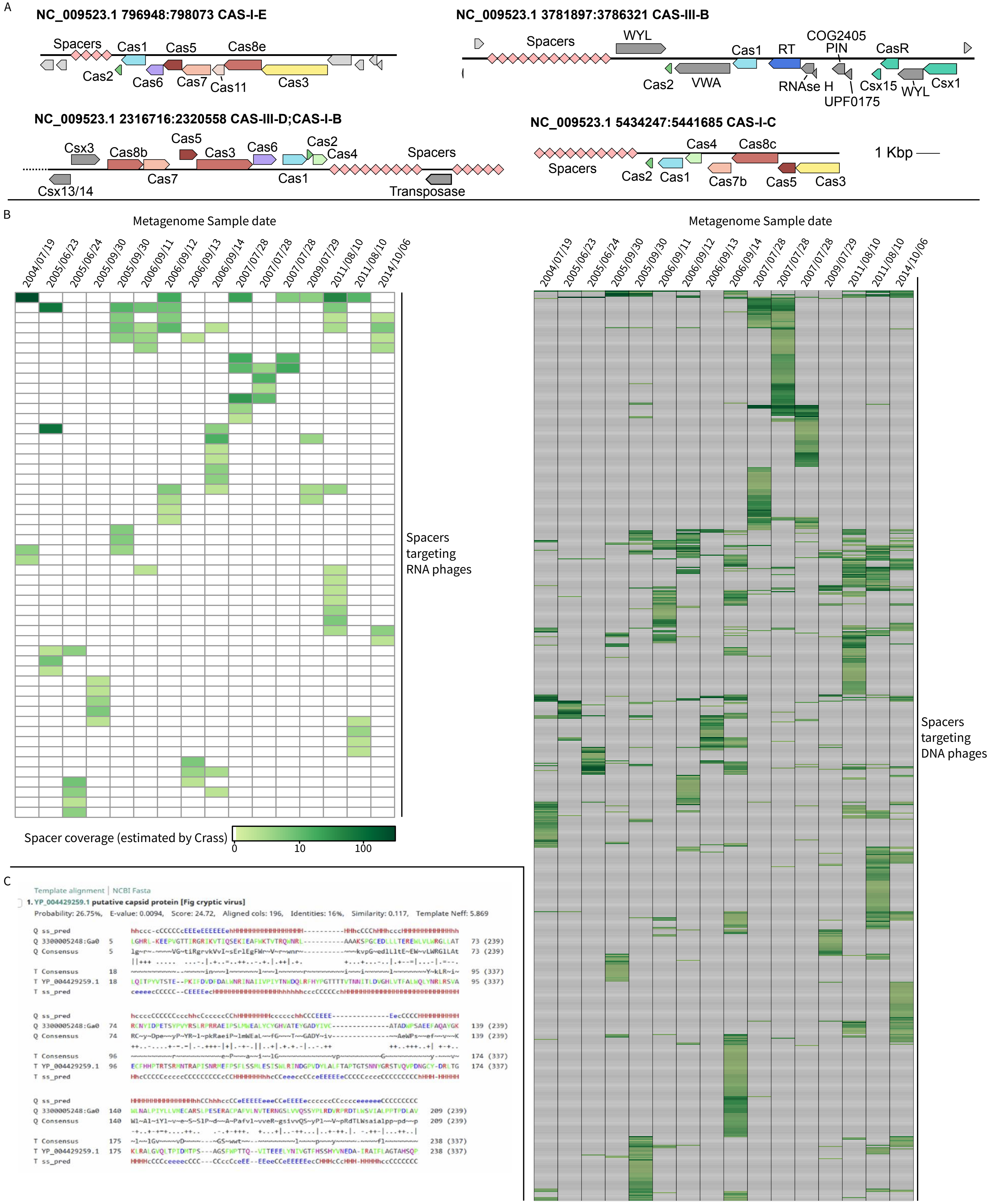

### Fig. S4 - Dataset types and associated virus types.pdf

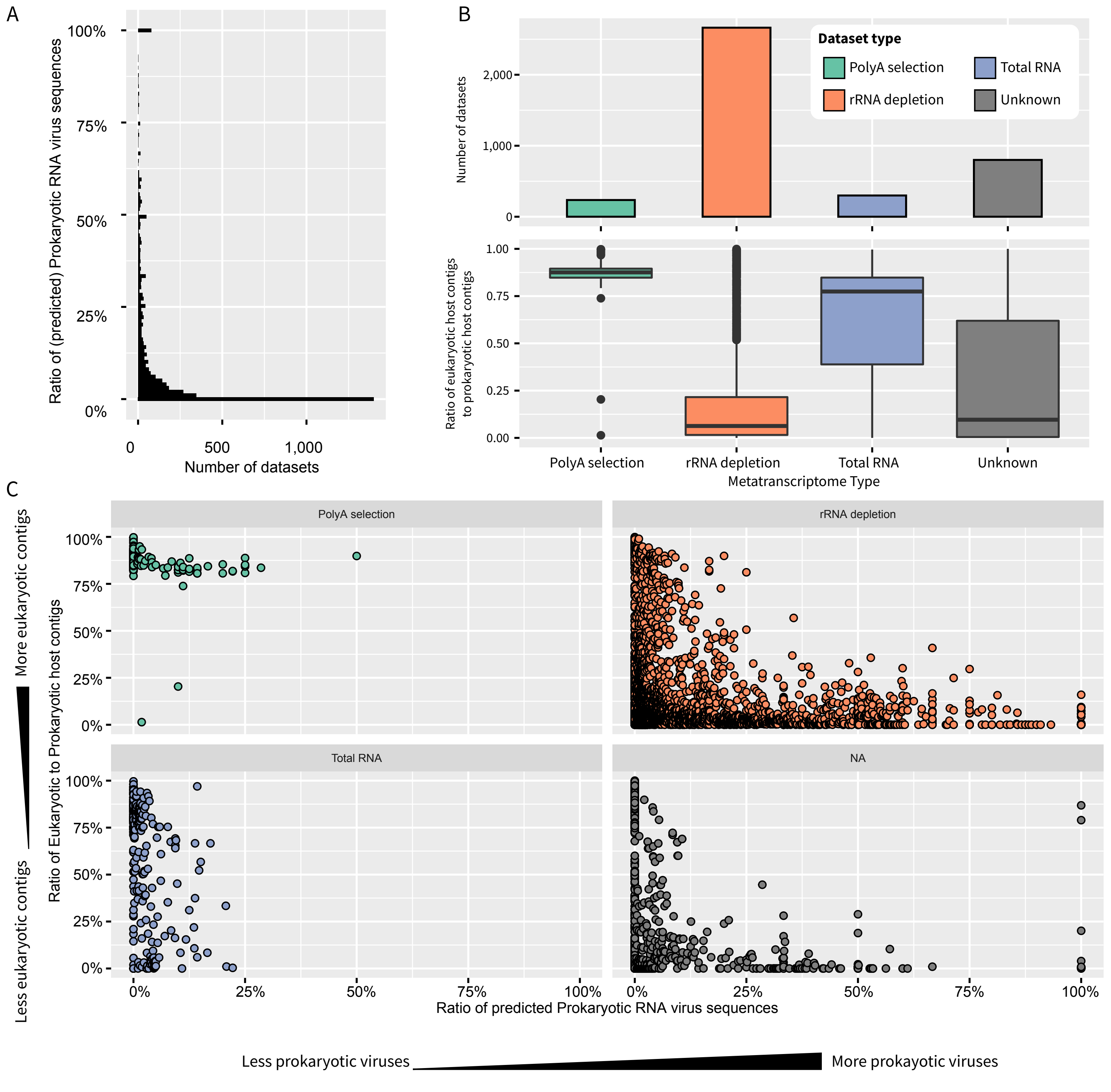

### Fig. S5 - Domain Segmentations_fusion_fission.pdf

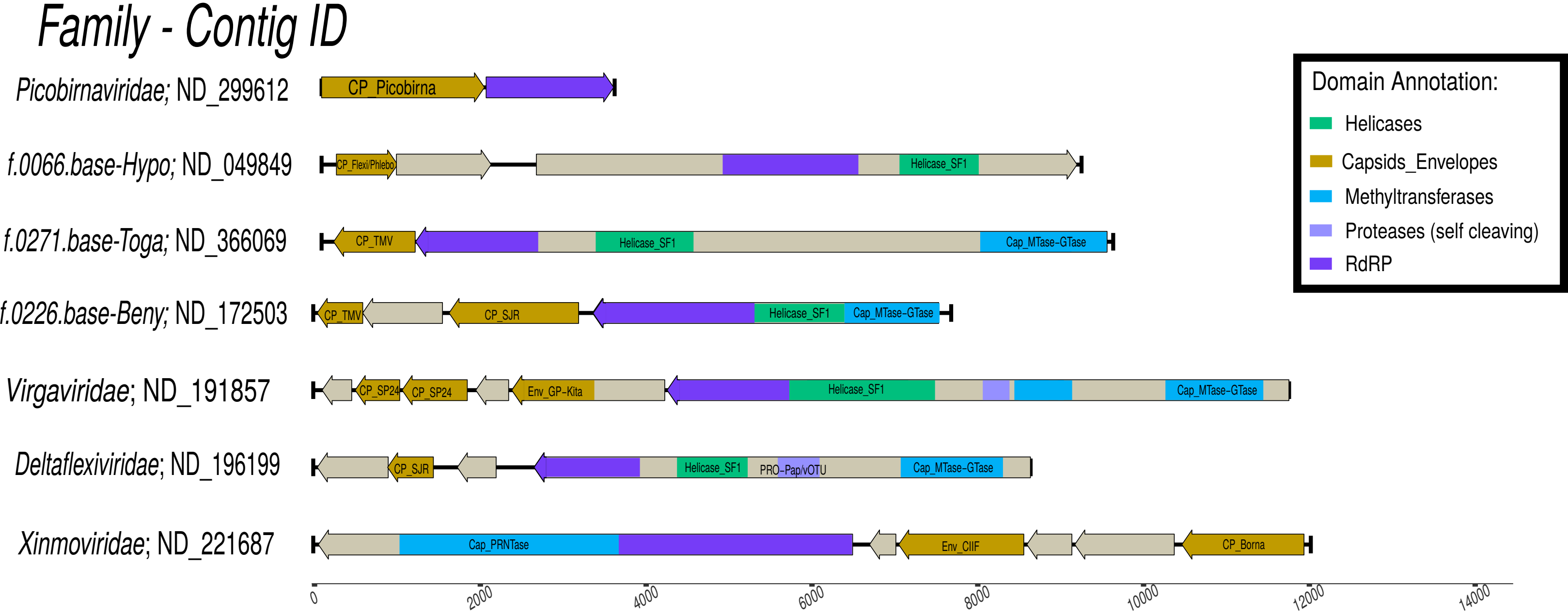

### Fig. S6 - Annotation_Pipeline.pdf

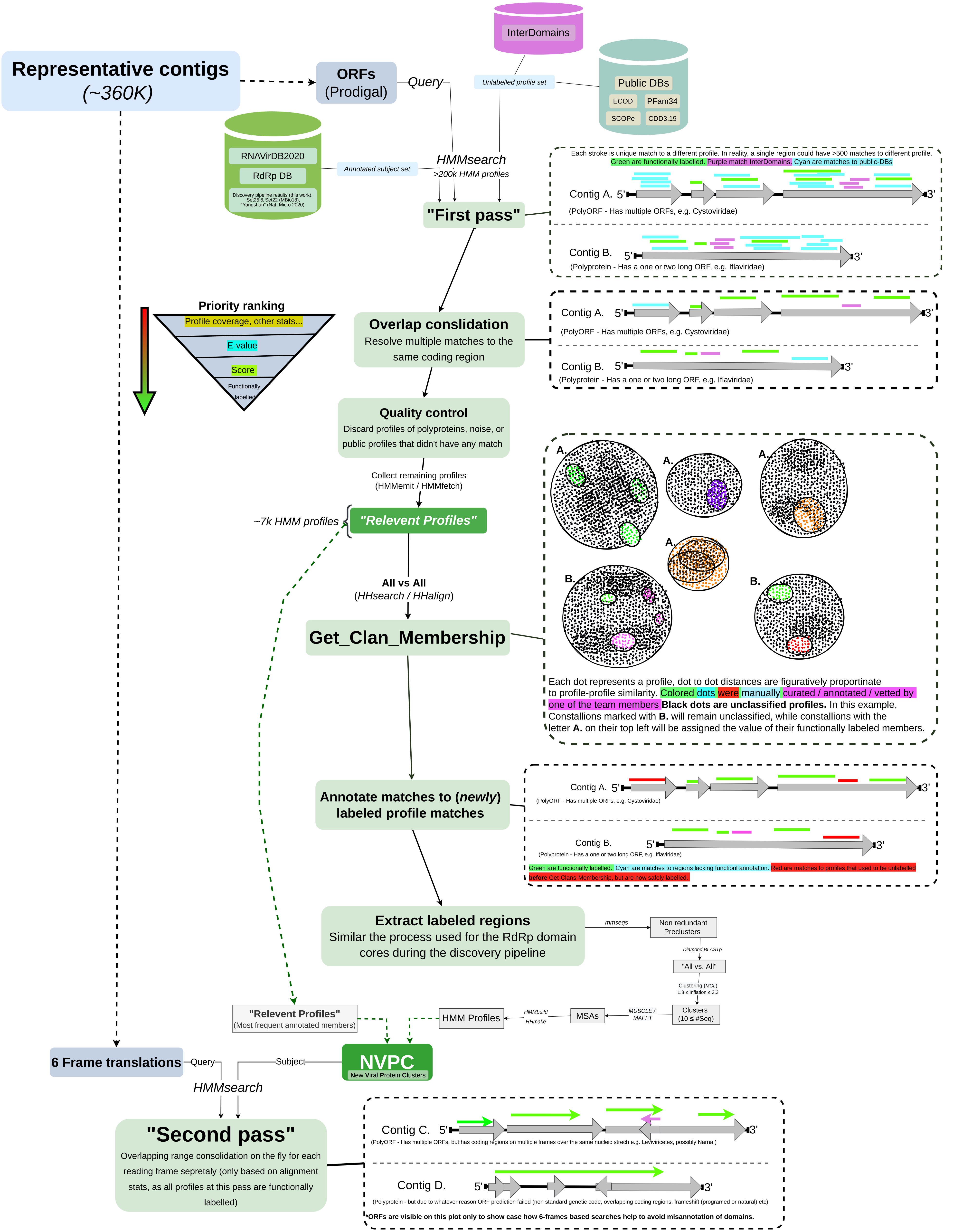

### Fig. S7- Identified Domain dist.pdf

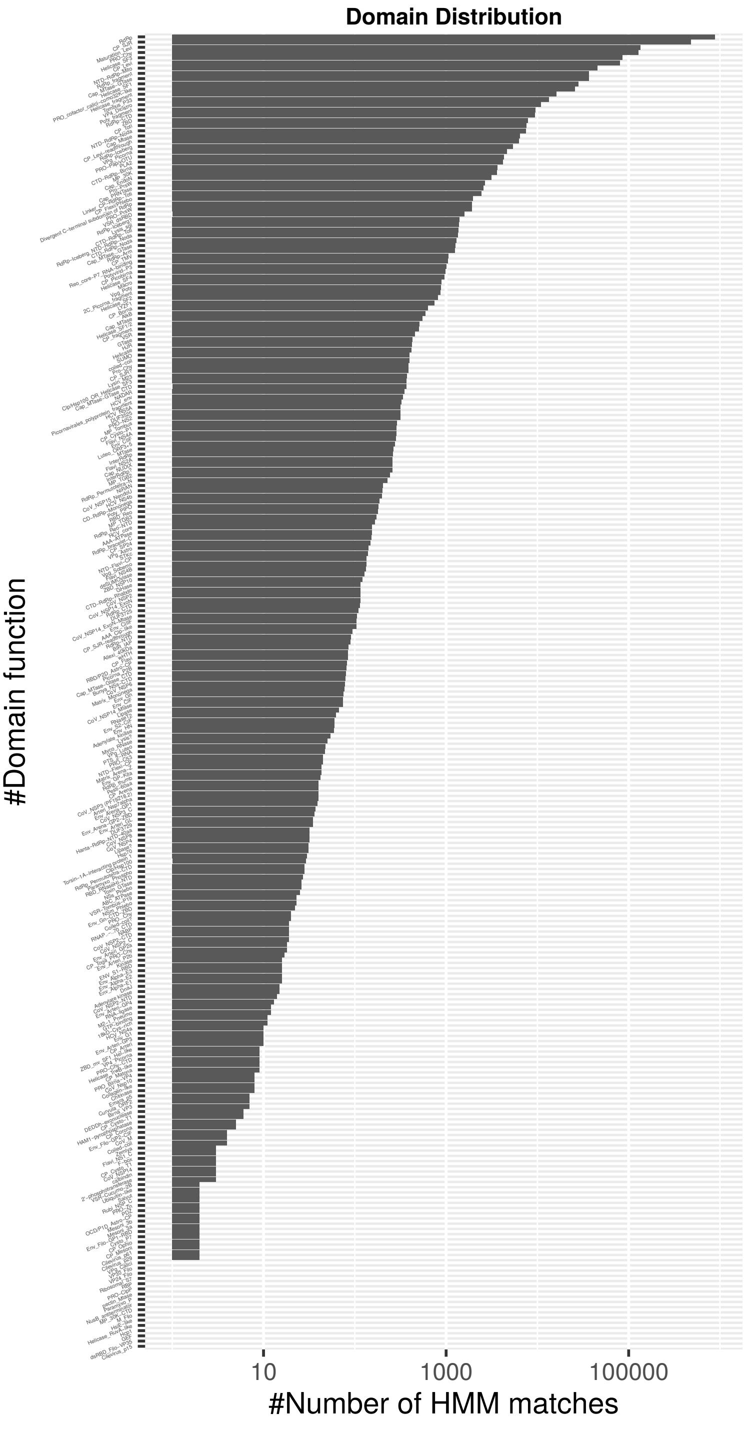
